# Progress toward the development of an effective vaccine for Extraintestinal pathogenic *E. coli* (ExPEC): The application of the multiple-protein subunits vaccine in different murine models

**DOI:** 10.1101/2023.05.31.543151

**Authors:** Yikun Xing, Justin R. Clark, James D. Chang, Jacob J. Zulk, Dylan M. Chirman, Felipe-Andres Piedra, Kathryn A. Patras, Anthony W. Maresso

**Affiliations:** Department of Molecular Virology and Microbiology, Baylor College of Medicine, Houston, Texas, United States of America; AILOR Labs, Vaccine Development Group, Baylor College of Medicine, Houston, Texas, United States of America; Alkek Center for Metagenomics and Microbiome Research, Baylor College of Medicine, Houston, Texas, United States of America

**Author notes:** **Author Contributions** **Conceptualization:** Yikun Xing, Anthony W. Maresso. **Data Curation:** Justin R. Clark. **Methodology:** Jacob J. Zulk, Dylan M. Chirman, Kathryn A. Patras. **Software:** Justin R. Clark, James D. Chang. **Formal Analysis:** James D. Chang. **Investigation:** Yikun Xing, Anthony W. Maresso. **Funding Acquisition:** Anthony W. Maresso. **Writing – original draft:** Yikun Xing. **Writing – review & editing:** Yikun Xing, Justin R. Clark, James D. Chang, Kathryn A. Patras, Anthony W. Maresso.

## Abstract

Extraintestinal pathogenic *E. coli* (ExPEC) is the primary Gram-negative bacterial pathogen, and the leading cause of life-threatening sepsis and urinary tract infections (UTI) in adults. The emergence and increasing prevalence of multidrug-resistance (MDR) ExPEC strains have led to considerable treatment failures, increased hospitalization rates, morbidity, and mortality. A prophylactic vaccine against ExPEC has the potential to reduce severe infection-related morbidity and mortality, helping to address the escalating antimicrobial resistance (AMR) crisis worldwide. The α-hemolysin (HlyA) is a critical, frequently detected secreted cytotoxic virulence factor in ExPEC, with HlyA-expressing ExPEC strains correlating with increased severity and infection dissemination in clinical levels. In this study, we assessed the protective efficacy of pro-HlyA (the inactive and immature precursor of HlyA) and Dual-Hit (a combination of pro-HlyA and SinH-3, the previously reported immunoglobulin-like domain-3 of the invasin-like autotransporter protein SinH), as ExPEC vaccine candidates. We demonstrated that immunizing mice with pro-HlyA or Dual-Hit significantly reduced bacterial burden and increased survival rates against pandemic ExPEC sequence type strains, ST73 (CFT073) and ST95 (UTI89), in the model of bacteremia and mortality. Both pro-HlyA or Dual-Hit immunizations also provided significant protection against UTI89 colonization in the bladder in the murine UTI model. Furthermore, vaccination with Dual-Hit provided enduring and robust protection against a mixture of ten typical high-virulent sequence types of ExPEC strains, resulting in a promising broad-spectrum vaccine candidate. These findings suggest that pro-HlyA and Dual-Hit might serve as highly effective vaccine targets and highlight the potential of these vaccine candidates for further development and evaluation.

## Introduction

Extraintestinal pathogenic *E. coli* (ExPEC) is the most common Gram-negative bacterial pathogen, causing a diverse range of clinical diseases at non-intestinal sites affecting all age groups [16]. In the flora of humans, the ExPEC group comprises the following major variants, including uropathogenic *E. coli* (UPEC), neonatal meningitis *E. coli* (NMEC), and septicemia associated isolates (SEPEC) [18]. ExPEC is the primary cause of bacteremia and urinary tract infections (UTIs), and a frequent cause of neonatal meningitis [17,19]. In the United States, over 970,000 sepsis cases are admitted annually, with an 8.7% yearly increase in incidence among hospitalized patients, accounting for over 50% of hospital deaths [25,26]. Based on the Centers for Disease Control and Prevention (CDC) multiple cause-of-death data (1999-2014), 6% of all deaths involved sepsis, 22% of these cases listing sepsis as the underlying cause [20]. Moreover, in 2017, approximately 48.9 million new cases of sepsis were recorded globally, with 11 million sepsis-related deaths were reported, accounting for 19.7% of all worldwide deaths [22]. In addition, sepsis management remains a major challenge for healthcare systems worldwide, resulting in a disproportionately high burden in terms of cost and hospital resource utilization. In the United States, sepsis management costs surpass those for any other disease, exceeding $24 billion in 2013, representing 13% of total hospital expenses and growing at three times the rate of other admissions [24].

The clinical management of pathogenic *E. coli* infections presents significant challenges due to the emergence and rising prevalence of multidrug-resistance (MDR) ExPEC strains. These strains frequently result in treatment failure, increased hospitalization rates, exacerbated morbidity and mortality, and are a primary driver of the global antimicrobial resistance (AMR) crisis. A recent review assessing the global burden of bacterial AMR across 204 countries and territories identified antibiotic-resistant pathogenic *E. coli* as a primary cause of mortality associated with drug resistance, accounting for approximately 200,000 deaths due to antimicrobial-resistant *E. coli* and around 800,000 deaths linked to AMR *E. coli* in 2019 [23]. The CDC reports that over two million people in the United States contract antibiotic-resistant diseases annually, with AMR contributing an additional $20 billion to direct healthcare costs and roughly $35 billion in lost productivity each year [27]. Furthermore, other studies found that antimicrobial-resistant ExPEC infections impair the capacity of human immune systems to combat infectious diseases, leading to various infectious complications in patients following prostate biopsy [28], solid organ transplant [29,30], or undergoing chemotherapy, dialysis, surgery, and joint replacement [27].

A single sequence type (ST) clonal group, known as ST131, belongs to the phylogenetic group B2, which is commonly associated with extraintestinal infections. ST131 is a clinically significant and globally dispersed pathogenic MDR *E. coli* lineage, which frequently produces extended-spectrum β-lactamases (ESBLs), often linked to CTX-M-15, and exhibits near universal resistance to fluoroquinolones [31]. Furthermore, ST131 *E. coli* isolates maintain a balance between colonization, virulence, and antibiotic resistance without incurring a fitness cost, attributed to their distinct virulence profiles and expanded number of virulence genes compared to non-ST131 isolates [32,33]. In the United States, ST131 significantly contributes to the resistance of clinical *E. coli* isolates, accounting for approximately 70% of fluoroquinolone resistant strains and over 50% of MDR isolates [34].

While ExPEC strains in the ST95 and ST73 lineages display a relatively lower incidence of antimicrobial resistance compared to ST131 lineages, they remain prevalent and well-recognized as highly pathogenic ExPEC strains among clinical isolates in patients with UTIs and bloodstream infections (BSIs) [35,39]. For instance, the ST95 lineage emerged as the most common dominant clonal group among *E. coli* isolates from patients with UTIs in California between 1999-2000 and 2016-2017 and ranked as the second most common clonal ExPEC group in a San Francisco study of patients with BSIs [36–38]. Meanwhile, the ST73 lineage was the most prevalent ST among major clonal ExPEC groups from urine and blood in 2000 in the United Kingdom and the third most common ST causing BSIs among patients admitted to a hospital in San Francisco [36–38].

A vaccine against ExPEC offers a prophylactic alternative to reduce mortality associated with severe *E. coli* infections and mitigate the AMR crisis. Over the past few decades, numerous groups have pursued protective immunity against ExPEC through various *E. coli* vaccine strategies. Some have focused on inactivated bacteria vaccines [41,42] or bacterial lysate vaccines [40,43]. However, the ineffectiveness of heat-killed inactivated bacterial vaccines in preventing uncomplicated UTIs and significant adverse effects of bacterial-lysate vaccines limited their prevention function [44]. Several attempts have been made to develop O-specific polysaccharide (O-antigen) conjugate vaccines [45–48]. For instance, ExPEC4V (bioconjugate O1A, O2, O6A, and O25B) has elicited functional antibody responses and is currently in phase 2 randomized controlled trial [49–53]. ExPEC9V, a newer vaccine in phase 3, targets invasive ExPEC in individuals aged 60 years and older with a history of UTIs [54]. However, the high heterogeneity of O-specific polysaccharides has limited the development of a polysaccharide vaccine capable of preventing ExPEC infections.

Alternative strategies include fimbrial-based vaccines such as FliC (or pilin) [61–63] and FimH (from type 1 fimbriae) [59,60], which promote antibody immune responses and prevent the colonization of various UPEC strains *in vivo*. Non-fimbrial-based vaccines, such as adhesin FdeC [58], PapG fimbrial adhesin [56], and Dr fimbriae [55], have also demonstrated protection against UTIs *in vivo* [57]. Moreover, four defined *E. coli* iron acquisition antigens (IroN, lutA, IreA, and FyuA) [66–69] and siderophores (iron-chelating compounds) [70] have generated strong immune responses, showing potential as ExPEC vaccine candidates [64,65]. Similarity, some toxin-based vaccine attempts, such as insoluble α-hemolysin [71,72], and auto-transporter toxin Vat [73], have exhibited some protection against ExPEC infection *in vivo*. Despite numerous anti-ExPEC vaccine investigations spanning over a couple of decades, no ExPEC vaccine has been approved by the U.S. Food and Drug Administration (FDA).

The α-hemolysin (HlyA) is a critical and commonly detected secreted cytotoxic virulence factor in ExPEC, associated with upper UTIs such as cystitis or pyelonephritis [81]. HlyA, a pore forming toxin belonging to the RTX toxin family (repeats in toxin), exhibits cytotoxic activity against various species and cell types, potentially causing severe tissue damage [78]. Additionally, HlyA can lyse erythrocytes and damage effector immune cells at high concentrations [76,79], promote bladder epithelial cell exfoliation, and induce apoptosis in target host cells at low concentrations [77]. Clinically, HlyA is linked to severe UTIs that can lead to renal complications and permanent renal scarring [75,80], and may also cause endothelial damage and renal vasoconstriction [82]. The *hlyA* gene exhibits high prevalence in clinical *E. coli* isolates from patients with bloodstream infections [83–85], UTIs [86–88], and pregnant women, reaching up to 61.5% in *E. coli* strains causing cystitis and 78.6% in strains causing pyelonephritis [89]. Furthermore, an analysis of the *hlyA* gene distribution among major ExPEC clones revealed its prevalence was significantly higher in ST73 (64.6%) compared to ST131 (14.8%) and ST95 (13.5%) and was more frequently found in strains from phylogroup clades B and C [90,91].

Considering the significance of HlyA in ExPEC pathogenesis and the high prevalence of hlyA sequence in clinical E. coli isolates, we sought to investigate the potential of pro-HlyA, the inactive and immature precursor form of HlyA [4], as a vaccine candidate against ExPEC infections. Additionally, we combined pro-HlyA with a previously reported antigen, SinH-3, which is the immunoglobulin-like domain-3 of the invasin-like autotransporter protein SinH [5], to create the Dual-Hit vaccine. In this study, we present data suggesting that both pro-HlyA and Dual-Hit confer rapid, enduring, and robust protection against various sequence types of ExPEC strains in multiple murine infection models.

## Materials and Methods

### Ethics statement

All methods performed on mice adhered to the relevant guidelines and regulations set forth by “The Guide and Care and Use of Laboratory Animals” (National Institute of Health). The Animal Use Protocol number AN-5177 was approved by Baylor College of Medicine’s Institutional Animal Care and Use Committee.

### Bacterial strains and culture conditions

The *E. coli* strains utilized in this study were obtained from a single colony grown on the Lysogeny Broth (LB) plate (10 g/L tryptone, 0.5 g/L sodium chloride (NaCl), and 5 g/L yeast extract). Bacterial cultures were incubated at 37L after resuscitation from a frozen stock (−80 °C, 10% glycerol). The ExPEC ST131 strains used in the study, JJ1886, JJ1901, JJ2050, JJ2528, and JJ2547, were kindly provided by James R. Johnson [1]. Uropathogenic *E. coli* (UPEC) strains UTI89 (O18:K1:H7, ST95) [2] and CFT073 (O6:K2:H1; ST73) [3] were kindly provided by Kathryn Patras. *E. coli* strains W0008 (ST127-like), W0044 (ST405-like), W0128 (ST648-like) were isolated from the blood or feces of hospitalized patients with bacteremia. The number of colony-forming units (CFU) administered was determined by correlating the optical density (OD) at 600 nm to the number of colonies observed after plating.

### The *hlyA* sequence distribution and HlyA alignment

Complete *E. coli* genomes were obtained from NCBI’s RefSeq database [119] and sorted into phylogroups as previously reported [91]. The sorted genomes were then categorized by sequence types using the MLST software (https://github.com/tseemann/mlst") which uses the PubMLST databases (https://pubmlst.org/) [120]. The categorized genome database was then used as a custom BLAST (version 2.8.1) [121–123] database to search for hits to the hlyA, hlyB, hlyC, and hlyD genes from the hly operon of *E. coli* UTI89 genome (accession: CP000243.1). To design Figure 1A, the underlying phylogenetic tree diagram was created using the autoMLST software [124] in concatenated alignment mode with 1,000 UltraFast Bootstrap replicates using representative genomes from the 8 *E. coli* phylogroups. The representative phylogenetic diagram was then overlaid with pie charts created in GraphPad Prism version 9.5.0 using BLAST hit results. Finally, the figures were combined using Biorender. All software used default settings unless otherwise specified. Open reading frames that overlapped with BLAST hits (described above) were extracted and translated using the bacterial translation code (translation table 11) in Geneious 2023.1.1. Translated HlyA sequences were then aligned using Geneious Alignment Software with free end gaps and otherwise default settings after truncated ORFs were removed. The resulting alignment was sorted by a phylogenetic tree annotation that was added using Geneious Tree Builder with a Jukes-Cantor/Neyman distance model and 1,000 Bootstrap replicates to create a consensus tree.

**Fig 1.**
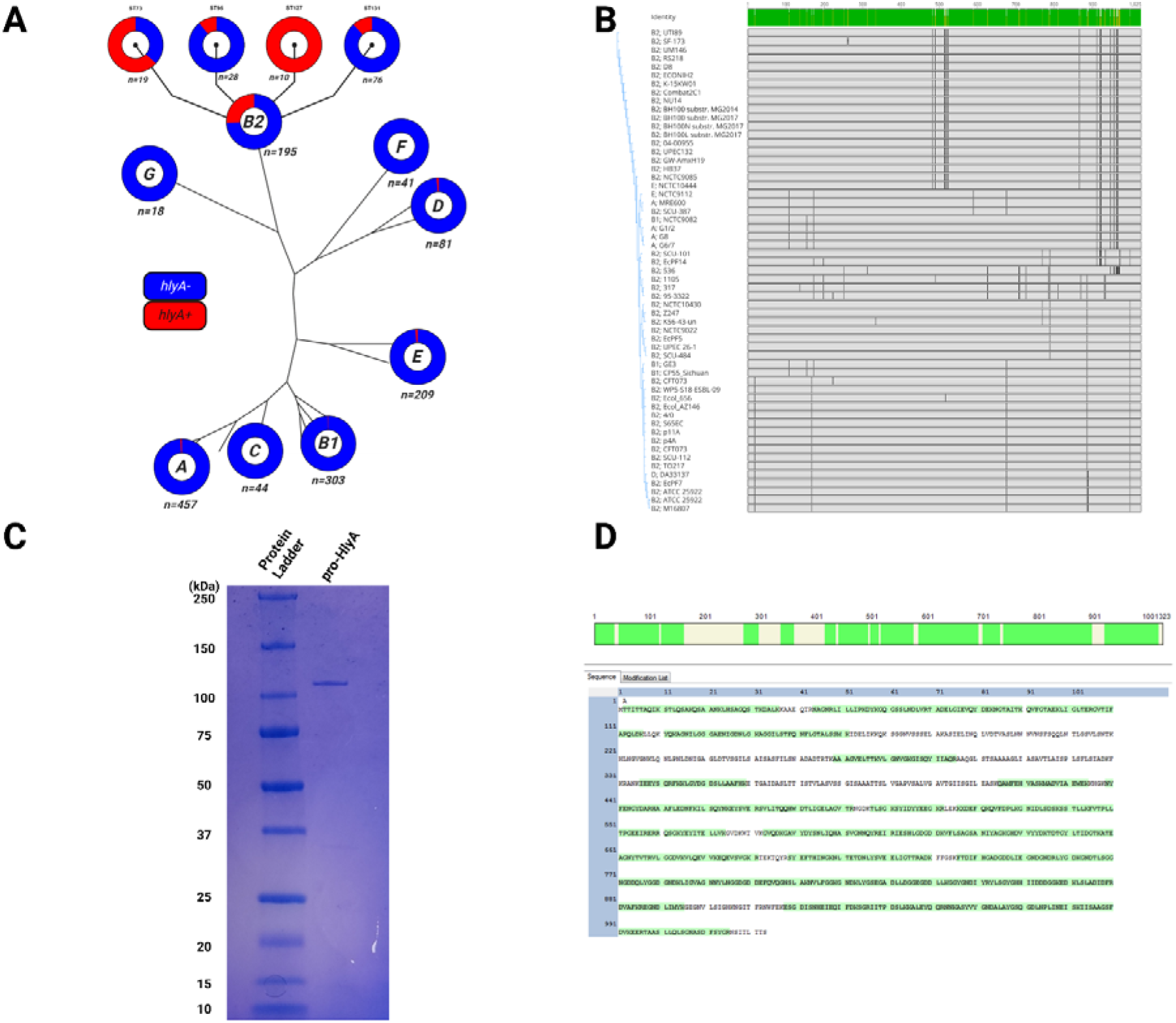
Hemolysin A is ExPEC-associated in genetical level. An analysis of a database of 1,348 complet *E. coli* genomes that have been phylogenetically categorized shows that the *hlyA* sequence is predominantly found in ExPEC-associated sequence types of the B2 phylogroup. **(A)** Phylogenetic representation of *hlyA* sequence distribution. BLAST was used to compare the *hlyA* nucleotide sequence to a database of complete *E. coli* genomes from NCBI’s Genbank that had been sorted into phylogroups using a previously described in-house method and into sequence types using MLST software (https://github.com/tseemann/mlst). Pie charts were made using GraphPad Prism, and the final figure wa created using Biorender. **(B)** Amino acid alignment of HlyA. Tickmarks represent disagreements with th majority consensus at that residue. Open reading frames overlapping with BLAST hits for *hlyA* were translated and aligned using Geneious Alignment and Geneious Tree Builder software (Geneious 2023.1.1). The tree annotation represents a consensus tree from 1,000 Bootstrap replicates. **(C)** Plasmid pSU-*hlyA* (encoding the *hlyA* sequence) and plasmid pK184-*hlyBD* (encoding *hlyB* and *hlyD* sequence) were co-transformed into *E. coli* BL21 (DE3) cells. Bacterial cultures expressing the recombinant pro-HlyA antigen secreted the protein into the supernatant, which was subsequently harvested, filtered, and concentrated. The purified antigen was analyzed by SDS-PAGE and stained with Coomassie blue stain buffer. Predicted size of pro-HlyA, 110kDa. The SDS-PAGE result was annotated using BioRender. **(D)** The coverage rate of pro-HlyA was determined by per-band sequencing through mass spectrometry.

### Prediction of Protein Structure for HlyA with AlphaFold2

The nucleotide sequence of HlyA was used to recreate the translated amino acid sequence using ExPASy. All six possible reading frames (three forward, three backward) were generated, and the frame that had the sequence for complete HlyA was used as the amino acid sequence for structure prediction. ColabFold’s AlphaFold2-Advanced Google Notebook (Google, Mountain View, CA) was used to generate predictions from amino acid sequences [108,109]. For multiple sequence alignment (MSA) necessary to build the consensus model for the structure of HlyA, we used MMseq2 [110,111]. Five prediction runs were run, with each run using a randomly chosen initiation point for the start of prediction runs. These models were ranked using the following two metrics: 1. pLDDT (predicted lDDT-Cα) with its ability to quantify the confidence of model per residue calculated by utilizing distances between Cα atoms in multiple reference models, and 2. AlphaFold-generated PAE (Predicted Aligned Error) for every residue, a numerical value of expected position error per residue [108,112]. The model with the highest average pLDDT and lowest PAE was chosen as the best-predicted structure of HlyA, and Amber Force Field was applied to relax the structure [113]. The predicted structure was compared against the list of previously solved structures of RTX toxins deposited on PDB aligning spatial coordinates of models by domains [114,115]. Additionally, Foldseek search was used to search for similar solved and AlphaFold-predicted structures on the AFDB-Swiss-Prot database through UniProt [116,117]. Additionally, these structures aligned with the predicted structure by UCSF ChimeraX’s alignment feature using the Needleman-Wunsch algorithm with BLOSUM-62 similarity matrix [118]. ChimeraX was used for analyzing the structural features of the predicted model, determining local physical properties within domains, and visualizing the model.

### Plasmid construction

The plasmid for the candidate vaccine antigen SinH-3 was constructed using a previously described method [5]. The plasmids pSU-*hlyA* (encoded the candidate vaccine antigen pro-HlyA, Uniprot entry: P08715), and pK184-*hlyBD* (encoded the necessary transport complex components HlyB and HlyD) were kindly provided by Lutz Schmitt [4]. The mRNA plasmid was constructed by cloning the SinH-3 gene from ExPEC sequence type 131 (ST131) strain JJ1887 genomic DNA (SinH-Ig-like domains-3, encoding the C-terminal passenger Ig-like domain-3 fragment of sinH, amino acid residues 602 to 724) and the pro-HlyA was cloned from *E. coli* (Uniprot entry: P08715). Both sequences (*hlyA* and *sinH-3*) were submitted to Creative Biolabs for constructing a *sinH-3*: *hlyA* mRNA construct with IL-2 signal peptide and P2A self cleaved sequence, called pIVTScrip-mRNA-IL2-sig_hlyA-P2A-IL2-sig_sinH (hereinafter named Dual-Hit mRNA construct). The Dual-Hit mRNA constructs were further enclosed in Cationic Lipid Nanoparticle (Cationic Lipid Nanoparticle (SM-102/DSPC/Cholesterol/DMG-PEG= 50: 10: 38.5: 1.5) and stored in Tris-based buffer at –80°C.

### Vaccine antigens preparation

The recombinant SinH-3 protein was expressed as fusions with glutathione-S-transferase (GST) using *E. coli* BL21(DE3) and purified as previously described [5]. To purify the recombinant protein pro-HlyA, the plasmid pSU-*hlyA*, containing the C-terminal secretion signal of HlyA (Uniprot entry: P08715), and pK184-*hlyBD*, encoding HlyB and HlyD essential for the transport complex, were co-transformed into *E. coli* BL21(DE3) cells [4]. A single pSU-*hlyA* and pK184-*hlyBD* co-transformed *E. coli* BL21(DE3) colony was used to inoculate a 300 ml baffled flask containing 150 ml of Lysogeny broth (LB) medium and cultured overnight. The overnight culture was then used to inoculate a 2 L baffled flask containing 800 ml of LB medium, which was grown at 37L until it reached optical density at 600 nm (OD600) of 0.4-0.6. Gene expression was induced with 1mM Isopropyl β-D-1-thiogalactopyranoside (IPTG) (Sigma Aldrich, St. Louis, MO), and the culture was incubated overnight at 37°C and 150rpm. Following induction, the supernatant containing secreted pro-HlyA protein was collected by centrifugation (Thermo Scientific, Sorvall RC 6+, SLA-3000 (Rotor), 10,000 × g for 30 min at 4°C) and filtered through the 0.22μm Vacuum Driven Sterile Filters (Sigma-Aldrich, St. Louis, MO). The filtered supernatant was subsequently concentrated to 1 ml using Amicon Ultra-15 Centrifugal Filter Units (Millipore Sigma, Burlington, MA) with a 100 kDa molecular-weight cut-off (MWCO). All antigens were analyzed by SDS-PAGE and Coomassie Brilliant Blue staining, and expression of both purified proteins was confirmed by mass spectrometry as previously described [5]. Both recombinant proteins were stored at -20°C until further use and handled at 4°C. The control group consisted of culturing a single untransformed *E. coli* BL21(DE3) colony only (hereinafter referred to as control supernatant). The resulting supernatant was collected and concentrated using the exact same procedure as that used for the purification of pro-HlyA above.

### Experimental Animals

Experimental animals used in this study were 6-8 weeks old BALB/cJ mice obtained from Jackson Laboratories (Bar Harbor, ME). They were provided with sterile food and water ad libitum and housed in filtered cages with 3-4 mice per cage. All experimental procedures performed on mice were approved in accordance with relevant guidelines and regulations from “The Guide and Care and Use of Laboratory Animals” (National Institute of Health) and were approved by Baylor College of Medicine’s Institutional Animal Care and Use Committee under protocol number AN-5177.

### Vaccination

For experiments involving protein-subunit vaccines, purified proteins were mixed with alum adjuvant (G-Bioscience, St. Louis, MO) in a 2:1 ratio of antigen to adjuvant, following the manufacturer’s guidelines. Female BALB/cJ mice (6-8 weeks old) were given three subcutaneous injections (S.C) of either 50 μg of pro-HlyA or a mixture of SinH-3 and pro-HlyA (50 μg each) on days 0, 14, and 28. Control groups were either given vaccinations control samples (comprising the same volume of control supernatant (30 μl), described above, mixed with alum adjuvant (30 μl) or left unvaccinated. In experiments involving mRNA vaccines, 6 week-old male BALB/cJ mice were given three intramuscular injections (I.M) of either 2 μg Dual-Hit mRNA construct (low-dose group, 40 μl to one hind leg muscle), 5 μg Dual-Hit mRNA construct (high-dose group, 40 μl to one hind leg muscle) [6], or 50 μl of Tris-based buffer (control group).

### Murine Model of Bacteremia (UTI89)

*E. coli* strains UTI89 were cultured under specified conditions one day prior to injection. On the day of injection (day 42), the strains were subcultured in LB broth at a ratio of 1:100 to an OD600 of approximately 0.6 (Log phase, ∼1 × 10^8^ CFU/ml), harvested by centrifugation (3,500 × g for 20 min at 4°C, Centrifuge 5702 R, Eppendorf North America, Framingham, MA), and suspended in an equivalent amount of 1 × PBS. Mice were intraperitoneally injected with 50 μl of the *E. coli* strain suspension (1 × 10^8^ CFU) on day 42, and the inoculum was quantified by plating dilutions onto LB agar. After 16-hours, mice were euthanized and necropsied to collect their kidney, spleen, and liver. The organs were homogenized in 1 ml 1× PBS using a BeadBlaster Refrigerated Homogenizer (Benchmark Scientific Inc, Sayreville, NJ, USA), and the organ homogenates were plated on LB agar plates and incubated at 37L to determine the number of bacteria or colony-forming units (CFU) per milliliter (mL).

### Murine Model of Mortality (UTI89 or CFT073)

On the day before injection, *E. coli* strains UTI89 and CFT073 were grown under the specified conditions. On the day of injection (day 42), mice were intraperitoneally injected with 50 μl of either UTI89 or CFT073 suspension (1 × 10^8^ CFU). Mice were closely monitored twice daily for ten days to observe morbidity and mortality. Survival data were collected over time, and moribund or dead animals were euthanized and necropsied to determine bacterial levels in their kidney, spleen, and liver. The organs were homogenized in 1 ml 1 × PBS using a BeadBlaster Refrigerated Homogenizer, and the organ homogenates were plated on LB agar plates and incubated at 37L to determine the number of bacteria or CFU per milliliter (mL). Moribundity was determined based on multiple observable features, including rough coat, hunched posture, lethargy, and hyperpnea.

### Murine Model of Urinary Tract Infection (UTI89 or CFT073)

UPEC strains UTI89 and CFT073 were grown and prepared as previously described. On day 42, mice were transurethrally inoculated with 50 μl of a UPEC strain suspension (1 × 10^8^ CFU). The inoculum was quantified by plating dilutions onto LB agar. After 72 hours, mice were euthanized and necropsied to collect bladders. The bladders were homogenized in 500 μl 1 × PBS using a BeadBlaster Refrigerated Homogenizer, and the organ homogenates were plated on LB agar plates and incubated at 37L to determine the number of bacteria or CFU per milliliter (mL).

### Murine Model of Mortality from Mixture of Ten Strains

Ten *E. coli* strains, representing typical sequence types (STs) of ExPEC, were grown and prepared as previously described. On day 42, mice were intraperitoneally injected with 50 μl of a mixture of ten ExPEC strains (equally mixed, a total of 1 × 10^8^ CFU). The inoculum was quantified by plating dilutions on LB agar. Mice were monitored twice daily for ten days to observe their survival. Survival data were collected over time, and moribund or dead mice were euthanized and necropsied to determine bacterial levels in their kidney, spleen, and liver. The organs were homogenized in 1 ml 1 × PBS using a BeadBlaster Refrigerated Homogenizer, and the homogenates were plated on LB agar plates and incubated at 37L to determine the number of bacteria with CFU per milliliter (mL). Moribundity was determined by observing multiple features, including rough coat, hunched posture, lethargy, and hyperpnea.

### Statistical analyses

Graphs and statistical analyses were conducted using Graphpad Prism version 9 (GraphPad Software, San Diego, CA). Significance was determined using the Mann-Whitney U test (two groups) or Kruskal-Wallis analysis of variance (ANOVA) with Dunn’s multiple comparisons correction (more than two groups). Survival curves were compared using the Genhan-Breslow Wilcoxon curve comparison. A 95% confidence interval was used for all statistical analyses, with alpha values of 0.05. Statistical significance was determined if calculated P-values were less than 0.05. The lines of all the bar graphs were at the median with 95% confidence intervals (CI). The statistical significance is represented as one star (*) for *P* <L0.05, two stars (**) for *P* <L0.01, three stars (***) for *P* <L0.001, and four stars (****) for *P* <L0.0001. The box-and whisker plots and Kaplan Meier survival curves were generated using Graphpad Prism 9 and annotated with BioRender.

## Results

### Hemolysin (HlyA) is genetically ExPEC-associated

Using a database of 1,348 complete *E. coli* genomes previously categorized by our lab, we were able to locate 65 genomes that contained BLAST hits for the *hlyA* sequence [91]. The phylogroup distribution of the *hlyA* sequence shows it is predominantly found in phylogroup B2, specifically the ExPEC and UPEC-associated sequence types (STs) ST73, ST95, ST127, and ST131 (Fig. 1A). Alignment and phylogenetic analysis of the amino acid sequences of HlyA suggests HlyA is highly conserved, with all alleles being >97% identical on the amino acid level (Fig. 1B). These results also suggest that the *hlyA* sequence has a horizontal pattern of transmission between phylogroups, as the alleles of hlyA within non-B2 phylogroups are nested within those of the B2 phylogroup (Fig. 1B).

### Candidate antigens expression and purification

To prepare the immunization, the candidate vaccine antigen pro-HlyA (Uniprot entry: P08715) was encoded into plasmid pSU-*hlyA*, while transport complex components HlyB and HlyD were encoded into plasmid pK184-*hlyBD*. Both plasmids were co-transformed into *E. coli* BL21 (DE3) cells. Bacterial cultures expressing recombinant pro-HlyA antigen secreted the protein into the supernatant, which was subsequently harvested, filtered, and concentrated. The recombinant proteins were visualized using sodium dodecyl sulfate-polyacrylamide gel electrophoresis (SDS-PAGE). Coomassie blue staining of the gels revealed a dominant band, presumed to be pro-HlyA (110 kDa) (Fig. 1C). The identity of the putative pro-HlyA protein was confirmed by mass spectrometry through per-band sequencing. Purified protein bands were resolved and digested in gel, and the tryptic peptides were analyzed on a nanospray LC-MS (liquid chromatography-mass spectrometry) system. The eluted peptides were directly electrosprayed into the mass spectrometer and analyzed by data-dependent acquisition (DDA). The coverage of the candidate antigen pro-HlyA, which is defined as the percentage of the protein sequence covered by identified peptides, was approximately 73%, with 97 peptides detected (Fig. 1D). This high sequence coverage substantiated the identity of the antigen, enabling its utilization in further experiments. The recombinant SinH-3 protein, fused to glutathione-S transferase (GST), was expressed in *E. coli* BL21(DE3) and purified following a previously described method (hereinafter named SinH-3) [5].

### Structural Prediction of HlyA with AlphaFold2

The AlphaFold2-predicted protein structure of HlyA reveals structural and organizational parallels with previously characterized RTX toxins. HlyA is predicted to comprise three domains: a putative N-terminal adenylate cyclase (residues 1-279, red), a three-helix bundle (residues 321-437, blue), a predominantly beta-helix C-terminal domain (residues 438-1023, green), and a linker connects the adenylate cyclase and helix-bundle domains (residues 280-320, grey) (Fig. 2A). The N-terminal adenylate cyclase domain is predicted to be primarily composed of ten alpha helices with four beta strands that form a sheet. This domain is predicted to adopt a globular conformation with an outer surface of charged residues and resultant hydrophilic surfaces (Fig. 2B). Although the putative catalytic site for adenylate cyclase activity could not be pinpointed solely from its structure, its amino acid sequence similarities with previously characterized RTX toxins strongly suggest cyclase catalytic activity. The linker region between the adenylate cyclase and three helix bundles (residues 280-320) contains five glycine residues (Gly 280, 283, 285, 298, 307; purple). These five residues would grant the linker region ability to assume shapes necessary for the proper placement of the adenylate cyclase domain with respect to the helix bundle, allowing optimum catalytic activity for the proper functioning of HlyA (Fig. 2C).

**Fig 2.**
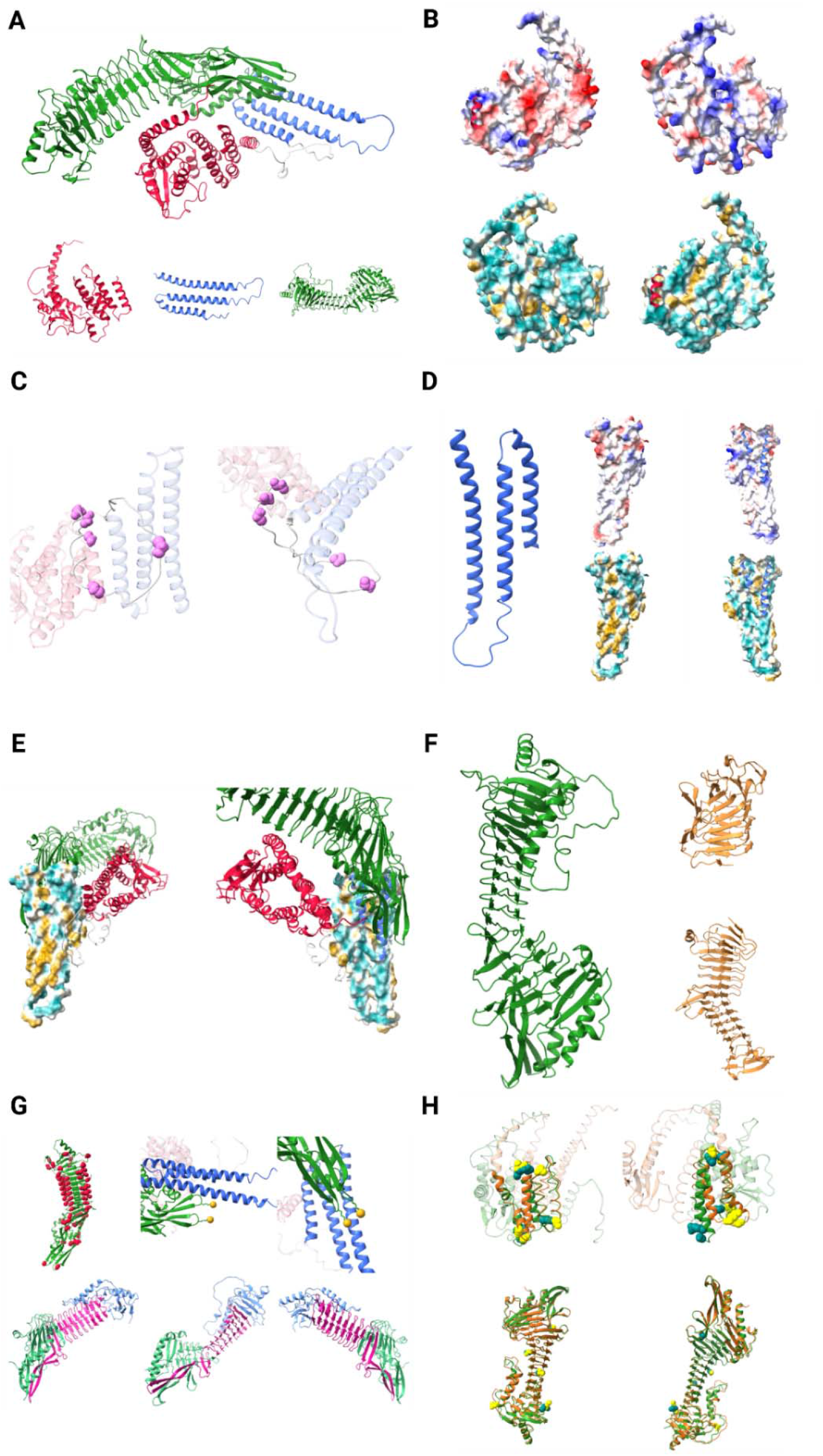
Predicting HlyA Structure Using AlphaFold2. **(A)** Overall predicted structure of HlyA. **(B)** Predicted structure of N-terminal adenylate cyclase domain. **(C)** The linker region between adenylate cyclase domain and three helix bundle. **(D)** Predicted structure of three helix bundle. **(E)** Hydrophobicity of three helix bundle in relation to other domains. **(F)** Predicted structure of C-terminal beta helix domain. **(G)** Organization of C-terminal beta helix domain. **(H)** Locations of residue differences between HlyA and P09983.

The next domain is predicted to assume three helices in alternating directions in a bundle (Fig. 2D). This prediction is based on the predicted alignment error plot for the model with the best prediction value, where amino acids that correspond to three helices in the linker region have low values of the predicted error to each other in all five prediction models (S1. Fig). Electrostatic surface maps of this region show predominantly uncharged residues along helices, with charged residues at ends of helices (blue: positively charged; red: negatively charged; white: uncharged). At both ends of helices, hydrophobicity maps show hydrophilic atoms; along the middle of all three helices, one face of helices is hydrophobic and the other hydrophilic (blue: hydrophilic; yellow: hydrophobic) (Fig. 2D). When these two surfaces are shown with rest of HlyA, the hydrophobic surface is distal to the C-terminal domain while hydrophilic surface proximal. This hydrophobic surface patch hints at the possibility of protein-protein interaction at this part of HlyA (Fig. 2E).

A predominant feature of the C-terminal domain is two-strand beta helix repeats that span the length of this domain (Fig. 2F). This domain shows high structural homology to several other well-characterized RTX toxins from bacterial pathogens, suggesting similar functions as the extracellular portion of the toxin that interacts with host surface proteins (hemolysin A from *P. mirabilis* (top, PDB: 4W8Q) and RTX fragment from *B. pertussis* AC toxin (bottom, PDB: 7RAH). Like these proteins, beta helix repeats feature glycine residues between beta strands that allow sharp changes in the direction of the strands (Fig. 2G). In addition to the beta helix that composes the majority of this domain, there is also a four-strand beta sheet predicted to be the next three helix bundle. In previous studies, acylation of two nitrogen atoms on lysine residues at two loops between strands by HlyC (Lys 563, 689) is known to be physiologically vital for HlyA to properly function as toxin, and side chains of these two lysines are situated right next to the three-helix bundle (above). The location of these two important lysine residues in the predicted model suggests correct modeling and placement of both helix bundle and C-terminal beta helix domains. As seen on other toxins with beta helices that are composed of multiple parts, there are three parts to the beta helix domain of HlyA (below). Intriguingly, the N-terminal part of the domain (green) inserts a beta strand into the sheet (residues 650-654). In addition, two lysines residues that require acylation for activity are composed of two different sections, with two strands from the middle domain (pink) inserted and places one of the two lysine at its proper location (residues 664-717) (Fig. 2G).

Hemolysin (P09983) from *E. coli* is the most structurally similar protein to this version of HlyA examined in the current study (Fig. 2H). When these two versions of HlyA are overlaid, and locations of amino acid differences are visualized, the most striking differences occur in the N terminal adenylate cyclase domain and C-terminal beta helix domain. In the N-terminal domain, differences that arise from structural variations can be found on turns that connect helices (green and teal: HlyA chain and different residues; brown and yellow: P09983 chain and different residues) (above). In the C-terminal domain, all amino acid residues with different compositions can be found on surfaces that are potentially involved in binding to either host or bacterial surfaces and possibly contribute to differential functioning by these two versions of HlyA (below) (Fig. 2H).

### Immunization with pro-HlyA antigen provides rapid protection against UTI89 infection in the murine model of bacteremia

UTI89, an ExPEC strain belonging to multilocus sequence type 95 (ST95) [7], has been isolated from patients with urinary tract infections and acute cystitis [8]. ST95, along with ST73 and ST131, is predominantly found in ExPEC strains and represents the second most prevalent clonal group in patients with bloodstream infections (BSIs) [9]. To evaluate the rapid protective efficacy of pro-HlyA in a UTI89 bacteremia model, experiment group mice were subcutaneously immunized with purified pro-HlyA combined with alum adjuvant (2:1 antigen/alum ratio), while the control group mice were injected with a mixture comprising equal volumes of control supernatant and alum adjuvant. On day 42, mice were followed by intraperitoneal injection of UTI89 (1 × 10^8^ CFU/mouse). The experimental vaccination scheme is shown in Figure 3A (Fig. 3A). After 16 hours of infection, mice were euthanized simultaneously, and their kidney, spleen, and liver were collected. The harvested organs were homogenized, and the bacterial burden of UTI89 in infected tissues was evaluated by quantifying colony-forming units (CFU) (Fig 3B - 3C).

**Fig 3.**
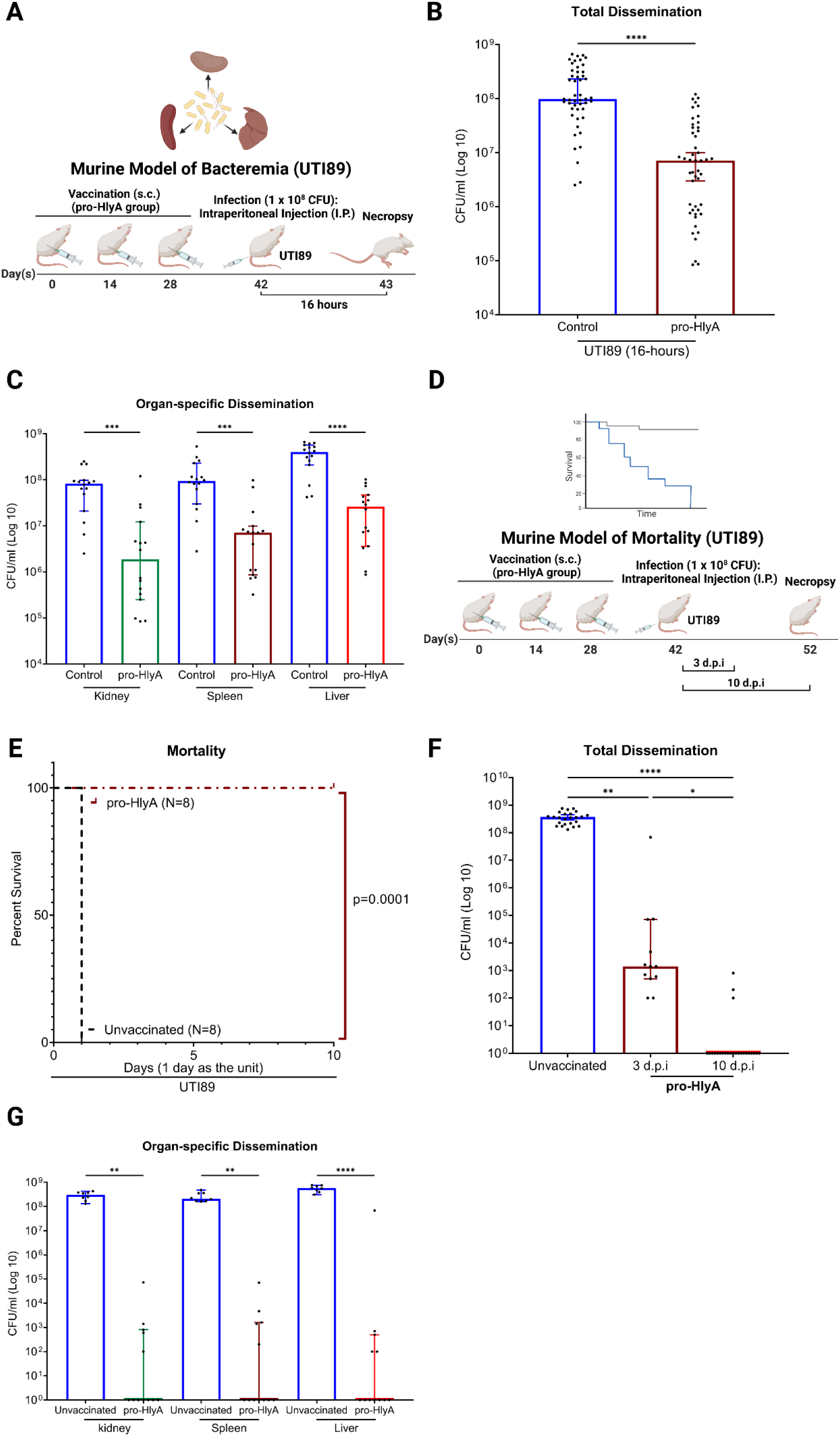
Evaluating the protective efficacy of pro-HlyA against UTI89 infections in the murine model of bacteremia and mortality. **(A)** The scheme of the murine bacteremia model using UTI89. Female BALB/cJ mice were subcutaneously immunized with either pro-HlyA (N=16) or control supernatant (N=16), followed by an intraperitoneal (I.P.) injection of 1 × 10^8^ CFU of UTI89. After 16 hours of infection, organs (kidney, spleen, liver) were harvested, homogenized, and plated to determine bacterial loads (CFU/ml). **(B)** Scatter plot with bar representing total UTI89 bacterial dissemination combining counts from all organs; **(C)** or the organ-specific UTI89 bacterial dissemination in each organ type post necropsy. **(D)** The scheme of the murine mortality model using UTI89. Female BALB/cJ mice were subcutaneously immunized with pro-HlyA (N=12) or left unvaccinated (N=8), followed by an intraperitoneal (I.P.) injection of 1 × 10^8^ CFU of UTI89. Mice were monitored twice daily for 10 days. The moribund or deceased mice were euthanized and necropsied to determine bacterial levels in organs (CFU/ml). Four surviving vaccinated mice were randomly chosen for euthanasia at 3 d.p.i, forming the 3 d.p.i group (N=4, excluded from survival rate calculation). All remaining survivors were euthanized at 10 d.p.i, forming the 10 d.p.i group (N=8). **(E)** The survival rate of pro-HlyA immunized mice after UTI89 infection was assessed using the Gehan-Breslow-Wilcoxon comparison. **(F)** Scatter plot with bar representing the total UTI89 bacterial dissemination combining counts from all organs at 3 d.p.i and 10 d.p.i; **(G)** or the organ-specific UTI89 bacterial dissemination in each organ type (combining counts from 3 d.p.i and 10 d.p.i) post-necropsy. Schemes were created in BioRender. Scatter plots with bars and Kaplan Meier survival curves were exported from Graphpad Prism 9 and annotated using BioRender.

Combining the counts from all cohorts (to assess the total effect of vaccination across all organs), mice immunized with pro-HlyA exhibited a significant reduction in bacterial burden across all organs (Adjusted *P*-value, *P* < 0.0001) (Fig. 3B). Compared to control group mice, pro-HlyA immunized mice showed a 1.14-log reduction in median UTI89 level at 16 hours post-infection, highlighting rapid and potent protection across multiple organs induced by pro-HlyA vaccination within a brief timeframe. Additionally, pro-HlyA vaccination led to significant reductions in bacterial levels within each organ type. Mice immunized with pro-HlyA showed a 1.65-log, 1.12-log, and 1.19-log reductions in median UTI89 levels in the kidneys, spleen, and liver, respectively (Adjusted *P*-value, kidney, *P* = 0.0006; spleen, *P* = 0.0008; liver, *P* < 0.0001), when compared to control group mice (Fig. 3C). These findings indicate the rapid, systemic protection against UTI89 bacteremia afforded by pro-HlyA immunization across multiple organs within a brief timeframe.

### Immunization with pro-HlyA increase survival rate in the UTI89 murine model of mortality

We subsequently assessed the long-term protective efficacy of pro-HlyA immunization. We investigated whether subcutaneous immunization with pro-HlyA increased the survival rate of vaccinated mice challenged with UTI89. The experimental group mice received three subcutaneous immunizations of pro-HlyA on day 0, 14, and 28, while mice in the control group remained unvaccinated. On day 42, mice were intraperitoneally injected with UTI89 (1 × 10^8^ CFU/mouse) and monitored twice daily for morbidity and mortality over the next 10 days. Survival was recorded over time, and moribund or deceased mice were euthanized and necropsied to assess bacterial loads in the kidneys, spleen, and liver. If no vaccinated mice died of infection, a small number of surviving vaccinated mice was randomly selected and euthanized at 3 days post-infection (d.p.i), forming the 3 d.p.i group (excluded from survival rate calculation), to assess changes in bacterial burden within infected organs over time. At 10 d.p.i, all remaining vaccinated mice were euthanized, forming the 10 d.p.i group. The harvested organs were homogenized, and UTI89 bacterial burden within infected tissues was quantified by determining CFU. The vaccination scheme used in this study is shown in Figure 3D (Fig. 3D).

Our results indicated that unvaccinated mice died within 1 d.p.i, whereas none of the pro-HlyA vaccinated mice died during the 10-day observation period, resulting in a 100% survival rate at 10 d.p.i (Adjusted *P*-value, *P* = 0.0001) (Fig. 3E). The bacterial burden results in pro-HlyA vaccinated mice correlated with survival rates. Combining the counts from all organs, pro-HlyA immunized mice exhibited a significant reduction in UTI89 bacterial burden at both 3 d.p.i (Adjusted *P*-value, *P* = 0.0060) and 10 d.p.i (Adjusted *P*-value, *P* < 0.0001), compared to unvaccinated mice. Moreover, in comparison to unvaccinated mice, the pro-HlyA vaccination resulted in approximately 5.42-log and 8.57-log reductions in median UTI89 bacterial burden at 3 d.p.i and 10 d.p.i, respectively (Fig. 3F). Furthermore, pro-HlyA vaccinated mice surviving for 10 d.p.i showed a 3.15-log significant reduction in median UTI89 levels (Adjusted *P*-value, *P* = 0.0147) compared to those euthanized at 3 d.p.i, suggesting enduring and consistent protection against UTI89 infection conferred by pro-HlyA immunization. Additionally, combining the counts at both 3 d.p.i and 10 d.p.i and analyzing by each organ, pro-HlyA immunized mice demonstrated a significant decrease in bacterial loads across multiple organs compared to unvaccinated mice (Adjusted *P*-value, kidney, *P* = 0.0013; spleen, *P* = 0.0040; liver, *P* < 0.0001) (Fig. 3G). This observation indicates a systemic reduction in bacterial levels due to pro-HlyA immunization, rather than localized effects.

### Immunization with pro-HlyA antigen alone provides insufficient protection against the mixture of ST131 ExPEC strains infection in the murine model of mortality

The ExPEC sequence type 131 (ST131) clone is the most common *E. coli* strain responsible for extraintestinal infections. In particular, the ST131-H30R lineage emerged clinically around the year 2000 and has become the leading antimicrobial-resistant and human clinical *E. coli* lineage in the United States [106,107]. Therefore, evaluating the protective efficacy of an ExPEC vaccine candidate against ST131 (including ST131-H30R lineage) is sorely required.

Here we evaluated whether pro-HlyA alone could provide sufficient cross-reactive protection against a mixture of five ST131 ExPEC strains that lacked the *hlyA* sequence (including ST131 H30R lineage) in the murine model of mortality. The experimental group mice received three subcutaneous immunizations with pro-HlyA with an alum adjuvant at a 2:1 ratio (antigen/alum) on days 0, 14, and 28, while the control group mice were injected with equal volumes of control supernatant and alum adjuvant. On day 42, all mice were intraperitoneally challenged with a mixture of five ST131 ExPEC strains (1 × 10^8^ CFU/mouse in total, JJ1886, JJ1901, JJ2050, JJ2528, JJ2547, in equal proportions) (Fig. 4A). Mice were closely monitored over the next 10 days for morbidity and mortality twice daily. Survival data were collected over time, and moribund or deceased mice were euthanized and necropsied to determine the bacterial loads in kidneys, spleen, and liver. The pro-HlyA vaccinated mice that died within 1 d.p.i, were grouped as the 1 d.p.i group, to assess whether the antibodies induced by pro-HlyA provide rapid cross-reactive protection against ST131 ExPEC strains. At 10 d.p.i, all the remaining vaccinated mice were euthanized, together with the mice that died at 6 d.p.i and 9 d.p.i, formed the 10 d.p.i group. Harvested organs were homogenized, and bacterial burdens within infected tissues were quantified by determining CFU. Our findings demonstrate that mice in the control group died within 1 d.p.i. Among the vaccinated mice during the 10-day observation period, 11 out of 16 pro-HlyA vaccinated mice died within 1 d.p.i, and two mice died at 6 d.p.i and 9 d.p.i, respectively, resulting in a survival rate of 18.8% at 10 d.p.i (Adjusted *P*-value, *P* = 0.0166) (Fig. 4B).

**Fig 4.**
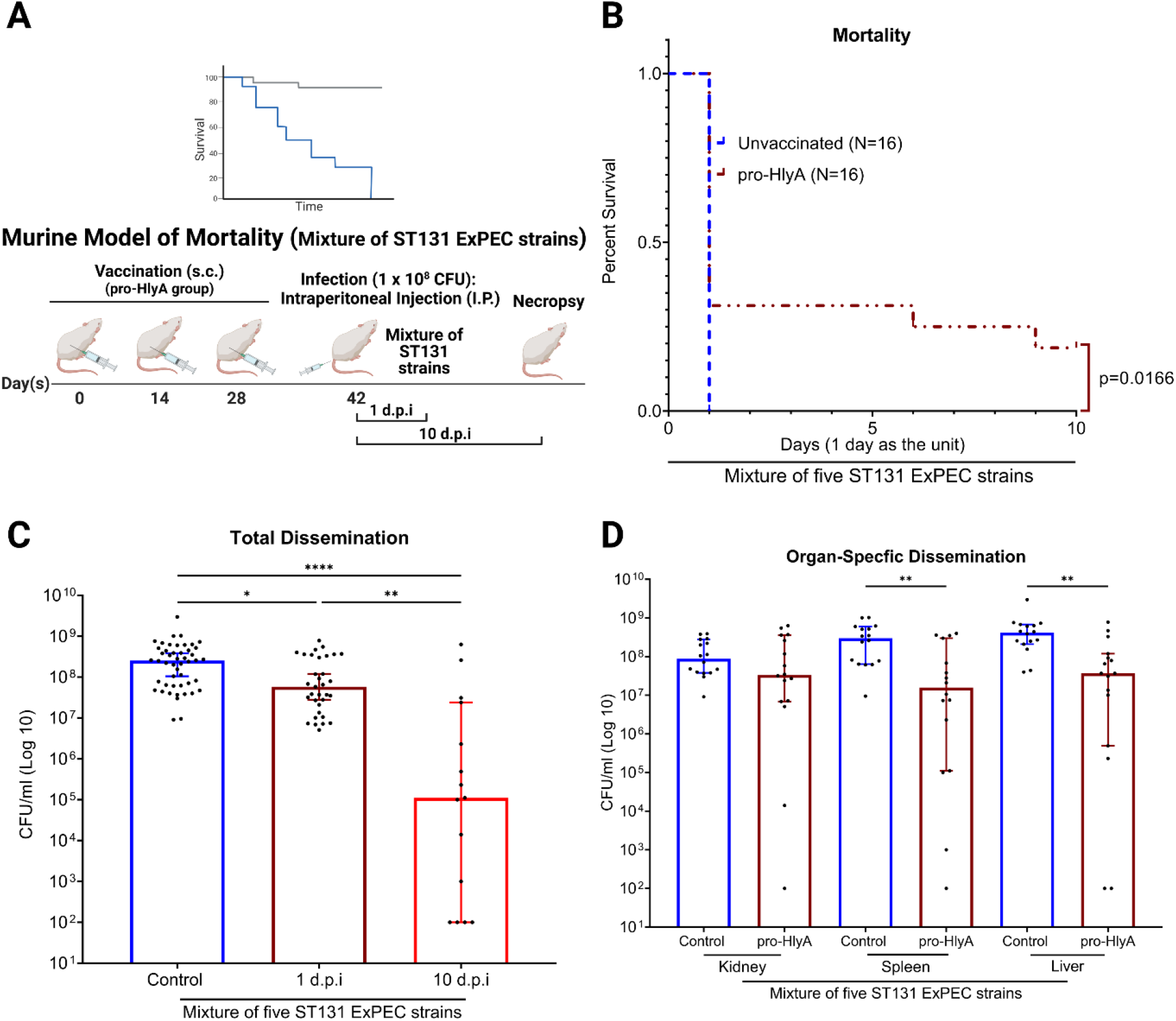
Evaluating the protective efficacy of pro-HlyA induced cross-reactive protection against a mixture of ST131 ExPEC strains in the murine model of mortality. **(A)** The scheme of the murine mortality model using a mixture of ST131 ExPEC strains. Female BALB/cJ mice were subcutaneousl immunized with pro-HlyA (N=16) or control supernatant (N=16), followed by an intraperitoneal (I.P.) injection of 1 × 10^8^ CFU of a mixture of ST131 ExPEC strains. Mice were monitored twice daily for 10 days. The moribund or deceased mice were euthanized and necropsied to determine bacterial levels in organs (CFU/ml). The pro-HlyA vaccinated mice that died within 1 d.p.i were grouped as the 1 d.p.i group (N=11). All the remaining vaccinated mice were euthanized at 10 d.p.i (N=3), together with the mice that died at 6 d.p.i and 9 d.p.i (N=2), forming the 10 d.p.i group (N=5). Organs (kidney, spleen, liver) were harvested, homogenized, and plated to determine bacterial loads (CFU/ml). **(B)** The survival rate of pro-HlyA immunized mice after a mixture of ST131 ExPEC strains infection was assessed using the Gehan-Breslow-Wilcoxon comparison. **(C)** Scatter plot with bar representing the total bacterial dissemination combining counts from all organs at 1 d.p.i and 10 d.p.i; **(D)** or the organ-specific bacterial dissemination in each organ type (combining counts from 1 d.p.i and 10 d.p.i) post-necropsy. Scheme were created in BioRender. Scatter plots with bars and Kaplan Meier survival curves were exported from Graphpad Prism 9 and annotated using BioRender.

When combining counts from all organs, pro-HlyA immunized mice exhibited a statistically significant reduction in bacterial burden at 1 d.p.i (Adjusted *P*-value, *P* = 0.0173) and 10 d.p.i (Adjusted *P*-value, *P* < 0.0001) (Fig. 4C) compared to the control group mice. Furthermore, upon combining bacterial burden counts at both 1 d.p.i and 10 d.p.i and analyzing them by each organ, pro-HlyA immunized mice only demonstrated statistically significant decreases in bacterial burdens in the spleen (Adjusted *P*-value, *P* = 0.0024) and liver (Adjusted *P*-value, *P* = 0.0012) compared to the control group mice (Fig. 4D). Therefore, these findings indicate that the cross-reactive protection induced by pro-HlyA vaccination offers rapid defense against ST131 ExPEC strains within a short time frame. However, the low survival rates observed in the vaccinated group suggest that this protection is insufficient to provide long-lasting effective protection in eliminating *E. coli* over time and to increase the survival rate after infection.

### Immunization with Dual-Hit confers protection against UTI89 infection in murine model of bacteremia

In a previous study, we demonstrated that SinH-3, a fragment corresponding to the immunoglobulin-like (Ig-like) domain-3 of the invasin-like autotransporter protein SinH, conferred robust systemic protection against infections caused by ST131 ExPEC strains in multiple murine models [5]. When challenged with a mixture of ST131 ExPEC strains that lack the *hlyA* sequence (including ST131-H30R lineage) in the murine model of mortality, given the insufficient protection induced by pro-HlyA immunization alone, we aimed to investigate the potential of a combination vaccine comprising SinH-3 and pro-HlyA as dual protein subunits (hereafter referred to as Dual-Hit) against several sequence types of ExPEC strains. We first assessed whether Dual-Hit maintained robust protective efficacy against representative ExPEC strains containing *hlyA* sequences, such as UTI89, in both bacteremia and mortality models.

To evaluate the rapid protective efficacy of Dual-Hit immunization in a UTI89 bacteremia model, experiment group mice were subcutaneously immunized with SinH-3 and pro-HlyA with an alum adjuvant at a 2:1 ratio (antigen/alum), while the control group mice were injected with a mixture comprising equal volumes of control supernatant and alum adjuvant. On day 42, mice received an intraperitoneal injection of UTI89 (1 × 10^8^ CFU/mouse). The experimental vaccination scheme is shown in Figure 5A (Fig. 5A). After 16 hours of infection, mice were euthanized simultaneously, and their kidneys, spleens, and livers were collected. The harvested organs were homogenized, and the bacterial burden of UTI89 within the infected tissues was quantified by measuring CFU (Fig 5B - 5C).

**Fig 5.**
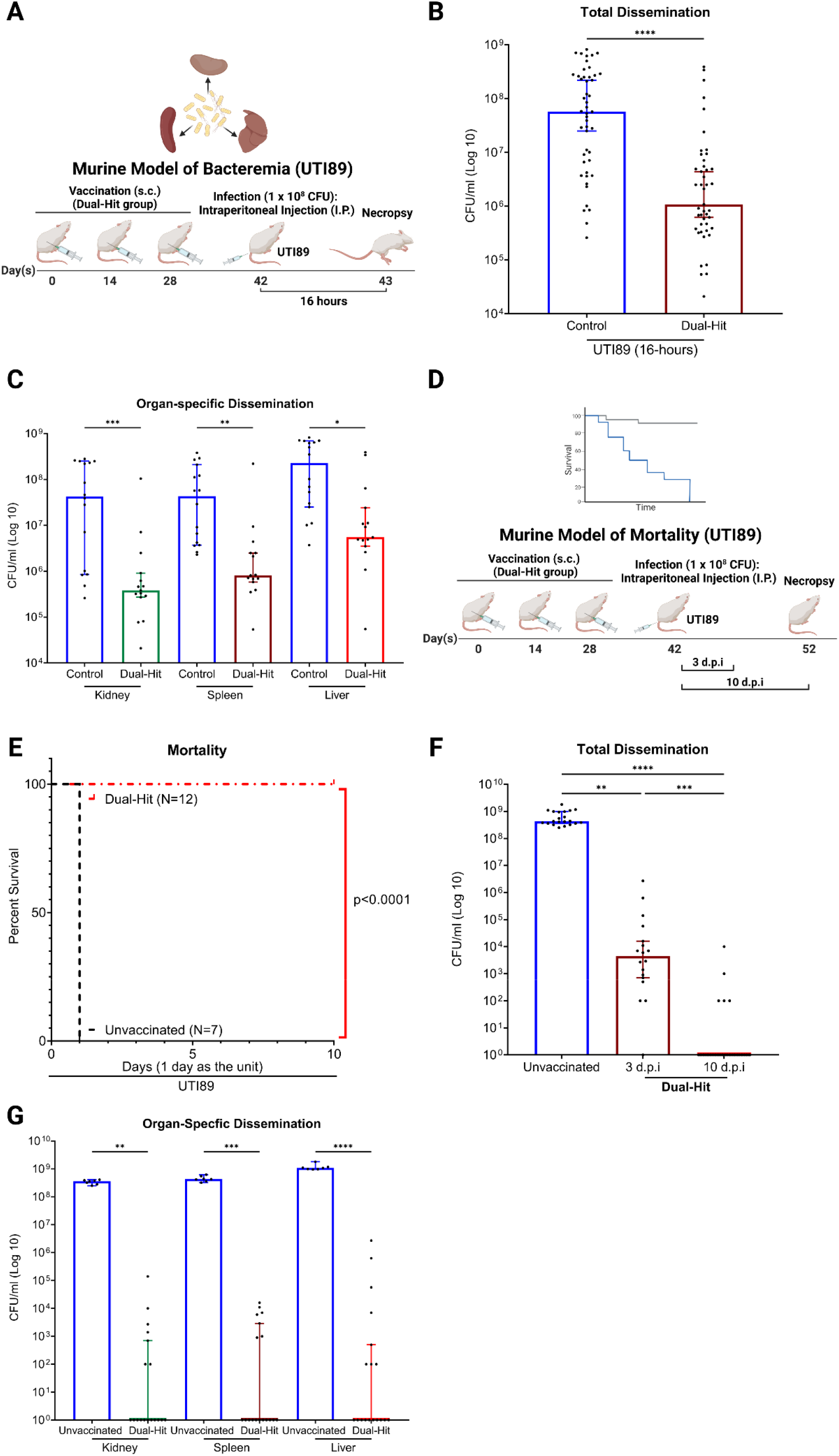
Evaluating the protective efficacy of Dual-Hit against UTI89 infections in the murine model of bacteremia and mortality. **(A)** The scheme of the murine bacteremia model using UTI89. Female BALB/cJ mice were subcutaneously immunized with either Dual-Hit (N=15) or control supernatant (N=16), followed by an intraperitoneal (I.P.) injection of 1 × 10^8^ CFU of UTI89. After 16 hours post infection, organs (kidney, spleen, liver) were harvested, homogenized, and plated to determine bacterial loads (CFU/ml). **(B)** Scatter plot with bar representing total UTI89 bacterial dissemination combining counts from all organs; **(C)** or the organ-specific UTI89 bacterial dissemination in each organ type post necropsy. **(D)** The scheme of the murine mortality model using UTI89. Female BALB/cJ mice were subcutaneously immunized with Dual-Hit (N=18) or left unvaccinated (N=7), followed by an intraperitoneal (I.P.) injection of 1 × 10^8^ CFU of UTI89. Mice were monitored twice daily for 10 days. The moribund or deceased mice were euthanized and necropsied to determine bacterial levels in organs (CFU/ml). Six surviving vaccinated mice were randomly chosen for euthanasia at 3 d.p.i, forming the 3 d.p.i group (N=6, excluded from survival rate calculation). All remaining survivors were euthanized at 10 d.p.i, forming the 10 d.p.i group (N=12). **(E)** The survival rate of Dual-Hit immunized mice after UTI89 infection was assessed using the Gehan-Breslow-Wilcoxon comparison. **(F)** Scatter plot with bar representing the total UTI89 bacterial dissemination combining counts from all organs at 3 d.p.i and 10 d.p.i; **(G)** or the organ-specific UTI89 bacterial dissemination in each organ type (combining counts from 3 d.p.i and 10 d.p.i) post-necropsy. Schemes were created in BioRender. Scatter plots with bars and Kaplan Meier survival curves were exported from Graphpad Prism 9 and annotated using BioRender.

Combining the counts from all cohorts, Dual-Hit vaccinated mice demonstrated a significant reduction in bacterial burden across all organs (Adjusted *P*-value, *P* < 0.0001) (Fig. 5B). Compared to the control group mice, Dual-Hit immunized mice exhibited an approximately 1.73-log reduction in median UTI89 level at 16 hours post-infection, indicating robust and rapid protection across multiple organs conferred by Dual-Hit immunization within a brief timeframe. Moreover, Dual-Hit vaccination resulted in significant reductions in bacterial levels within each collected organ. Compared to control group mice, those vaccinated with Dual-Hit exhibited approximately 2.05-log, 1.72-log, and 1.61-log reductions in median UTI89 levels in the kidneys, spleen, and liver, respectively (Adjusted *P*-value, kidney, *P* = 0.0009; spleen, *P* = 0.0047; liver, *P* = 0.0289) (Fig. 5C). These findings suggest that Dual-Hit immunization can still provide rapid, systemic protection against UTI89 bacteremia across multiple organs within a brief timeframe.

### Immunization with Dual-Hit increases survival rate in the UTI89 murine model of mortality

We subsequently assessed the protective efficacy of long-term protection conferred by Dual-Hit immunization. We investigated whether subcutaneous immunization with Dual-Hit increased the survival rate of vaccinated mice when challenged with UTI89. The experimental group mice received three subcutaneous immunizations of Dual-Hit on day 0, 14, and 28, while the control group mice remained unvaccinated. On day 42, mice were intraperitoneally challenged with UTI89 (1 × 10^8^ CFU/mouse). Over the following 10 days, mice were closely monitored twice daily for morbidity and mortality. Survival was recorded over time, and moribund or deceased mice were euthanized and necropsied to determine bacterial loads in the kidneys, spleen, and liver. If no vaccinated mice died of infection, a random small number of surviving vaccinated mice was selected and euthanized at 3 d.p.i, forming the 3 d.p.i group (excluded from survival rate calculation), to assess changes in bacterial burden within infected organs over time. At 10 d.p.i, all remaining vaccinated mice were euthanized, forming the 10 d.p.i group. Harvested organs were homogenized, and UTI89 bacterial burden within infected tissues was quantified by determining CFU. The vaccination scheme is shown in Figure 5D (Fig. 5D).

Our results showed that unvaccinated mice died within 1 d.p.i, whereas none of the Dual-Hit vaccinated mice died during the 10-day observation period, resulting in a survival rate of 100% at 10 d.p.i (Adjusted *P*-value, *P* < 0.0001) (Fig. 5E). Bacterial burden measurements in Dual-Hit vaccinated mice correlated with survival rates. When combining the counts from all organs, Dual-Hit immunized mice showed a significant reduction in UTI89 bacterial burdens at both 3 d.p.i (Adjusted *P*-value, *P* = 0.0037) and 10 d.p.i (Adjusted *P*-value, *P* < 0.0001), compared to unvaccinated mice. Moreover, in comparison to unvaccinated mice, the median UTI89 bacterial burden in Dual-Hit vaccinated mice was approximately 4.99-log and 8.63-log lower at 3 d.p.i and 10 d.p.i, respectively (Fig. 5F). Additionally, Dual-Hit vaccinated mice surviving for 10 d.p.i showed a 3.65-log significant reduction in median UTI89 levels (Adjusted *P*-value, *P* = 0.0003) compared to those euthanized at 3 d.p.i, indicating the persistent and sustained protection provided by Dual-Hit immunization against UTI89 infection. When combining bacterial burden counts at both 3 d.p.i and 10 d.p.i and analyzing by each organ, Dual-Hit immunized mice demonstrated significant reductions in bacterial burdens across multiple organs compared to unvaccinated mice (Adjusted *P*-value, kidney, *P* = 0.0013; spleen, *P* = 0.0003; liver, *P* < 0.0001) (Fig. 5G). These reduced bacterial levels were observed across all collected organs, suggesting a comprehensive reduction rather than localized effects provided by Dual-Hit immunization.

### Immunization with pro-HlyA or Dual-Hit confers partial protection against CFT073 and increases survival rates in the murine model of mortality

CFT073, a prototypical UPEC strain isolated from a female patient with acute pyelonephritis, belongs to phylogenetic group B2 and multilocus sequence type 73 (ST73) [10,11]. Notably, ST73 represents one of the most prevalent UPEC lineages, accounting for 11% and 16.6% of UPEC isolates obtained from UTI patients (including the elderly) in recent studies [12,13]. We investigated whether immunization with pro-HlyA or Dual-Hit confers robust protection against CFT073 in the murine model of mortality.

Mice received three subcutaneous immunizations with pro-HlyA or Dual-Hit on day 0, 14, and 28, while a control group remained unvaccinated. On day 42, mice were intraperitoneally injected with CFT073 (1 × 10^8^ CFU/mouse). Over the next 10 days, mice were monitored twice daily for morbidity and mortality. Survival was recorded over time, and moribund or deceased mice were euthanized and necropsied to assess bacterial loads in kidneys, spleen, and liver. Immunized mice that died or became moribund within 2 days formed 2 d.p.i group. At 10 d.p.i, all vaccinated remaining surviving mice were euthanized, forming the 10 d.p.i group. Harvested organs were homogenized, and CFT073 bacterial burden was quantified by determining CFU. The vaccination scheme is shown in Figure 6A (Fig. 6A). Our results indicated that all unvaccinated mice died within 1 d.p.i. whereas mice immunized with pro-HlyA exhibited a 25% survival rate at 10 d.p.i (Adjusted *P*-value, *P* = 0.0021), while Dual-Hit vaccinated mice showed a 42% survival rate at 10 d.p.i (Adjusted *P*-value, *P* = 0.0021) (Fig. 6B). Although some vaccinated mice died within 2 d.p.i, immunization with pro-HlyA or Dual-Hit significantly improved survival rates after CFT073 infections.

**Fig 6.**
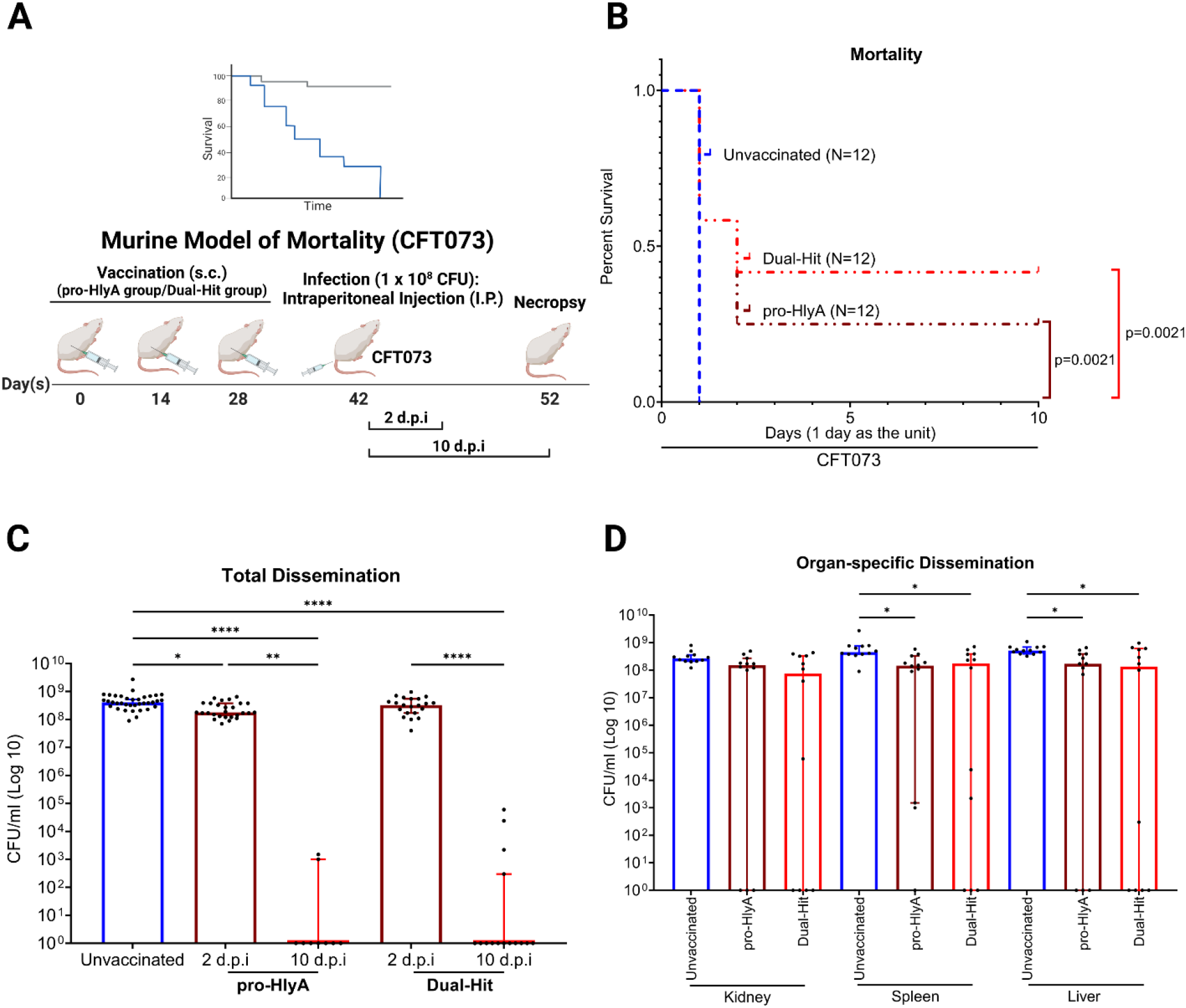
Assessing the protective efficacy of pro-HlyA and Dual-Hit against CFT073 infections in the murine model of mortality. **(A)** The scheme of the murine mortality model using CFT073. Female BALB/cJ mice were subcutaneously immunized with pro-HlyA (N=12), Dual-Hit (N=12), or left unvaccinated (N=12), followed by an intraperitoneal (I.P.) injection of 1 × 10^8^ CFU of CFT073. Mice were monitored twice daily for 10 days. The moribund or deceased mice were euthanized and necropsied to determine bacterial levels in organs (kidney, spleen, liver, CFU/ml). Vaccinated mice that died of infection within 2 d.p.i formed the 2 d.p.i group for each vaccinated cohort (pro-HlyA, N=9; Dual-Hit, N=7). All remaining survivors were euthanized at 10 d.p.i, forming the 10 d.p.i group for each vaccinated cohort (pro-HlyA, N=3; Dual-Hit, N=5). **(B)** Survival rates of pro-HlyA or Dual-Hit immunized mice following CFT073 infection were analyzed using the Gehan-Breslow-Wilcoxon comparison. **(C)** Scatter plot with bar representing the total CFT073 bacterial dissemination combining counts from all organs at 2 d.p.i and 10 d.p.i; **(D)** or the organ-specific CFT073 bacterial dissemination in each organ type (combining counts from 2 d.p.i and 10 d.p.i) post-necropsy. Schemes were created in BioRender. Scatter plots with bars and Kaplan Meier survival curves were exported from Graphpad Prism 9 and annotated using BioRender.

Bacterial loads in both pro-HlyA and Dual-Hit immunized mice correlated with survival rates. When combining counts from all organs, compared to unvaccinated mice, pro-HlyA immunized mice demonstrated a statistically significant reduction in CFT073 bacterial burden at both 2 d.p.i (Adjusted *P*-value, *P* = 0.0184) and 10 d.p.i (Adjusted *P*-value, *P* < 0.0001). Surviving pro-HlyA vaccinated mice exhibit an approximate 8.61-log reduction in the median level of CFT073 at 10 d.p.i relative to unvaccinated mice that died within 1 d.p.i. Additionally, pro-HlyA vaccinated mice surviving at 10 d.p.i showed a significant reduction in the median level of CFT073 compared to those moribund or died within 2 d.p.i (Adjusted *P*-value, *P* = 0.0035), indicating sustained and persistent protection caused by pro-HlyA immunization against CFT073 infection over time (Fig. 6C).

Similarly, surviving Dual-Hit vaccinated mice demonstrated a significant reduction in bacterial burden at 10 d.p.i compared to unvaccinated mice (Adjusted *P*-value, *P* < 0.0001), with an approximately 8.61-log reduction in the median level of CFT073 strain. However, there is no difference in bacterial burden between the 2 d.p.i Dual-Hit vaccinated group mice and unvaccinated group mice. Moreover, the median CFT073 levels were substantially reduced when comparing Dual-Hit vaccinated mice that survived at 10 d.p.i to those that died within 2 d.p.i (Adjusted *P*-value, *P* < 0.0001). These findings suggest that both pro-HlyA and Dual-Hit immunizations confer sustained and persistent protection against CFT073 infection (Fig. 6C).

Moreover, when combining bacterial burden counts at both 2 d.p.i and 10 d.p.i and analyzing them by each organ, pro-HlyA vaccinated mice exhibited a statistically significant reduction in bacterial burdens in the spleen (Adjusted *P*-value, *P* = 0.0146) and liver (Adjusted *P*-value, *P* = 0.0349) compared to the bacterial loads in unvaccinated mice (Fig. 6D). Similarly, Dual-Hit vaccinated mice showed a statistically significant reduction in bacterial levels in the spleen (Adjusted *P*-value, *P* = 0.0144) and liver (Adjusted *P*-value, *P* = 0.0186) compared to the bacterial burden in unvaccinated mice (Fig. 6D).

### Immunization with pro-HlyA or Dual-Hit confers protection against cystitis caused by UTI89 in the murine model of UTI

Urinary tract infections (UTIs) constitute a major global health concern, significantly contributing to morbidity in otherwise healthy females, with over 60% experiencing a diagnosis during their lifetime [14]. In the United States, the annual incidence of physician-diagnosed UTIs exceeds 10% for females and 3% for males. UPEC is the primary causative agent, accounting for approximately 80% of UTI cases [15]. Therefore, we evaluated the protective efficacy of pro HlyA or Dual-Hit against UPEC colonization in the bladder in the murine model of UTI. Female BALB/cJ mice were immunized subcutaneously with either pro-HlyA or Dual-Hit on days 0, 14, and 28, as previously described. On day 42, mice were transurethrally inoculated with 1 × 10^8^ CFU of typical UPEC strains (UTI89 or CFT073, Fig. 7A). After 72 hours of infection, bladders were harvested, homogenized, and bacterial loads of UTI89 and CFT073 were determined by quantifying CFUs.

**Fig 7.**
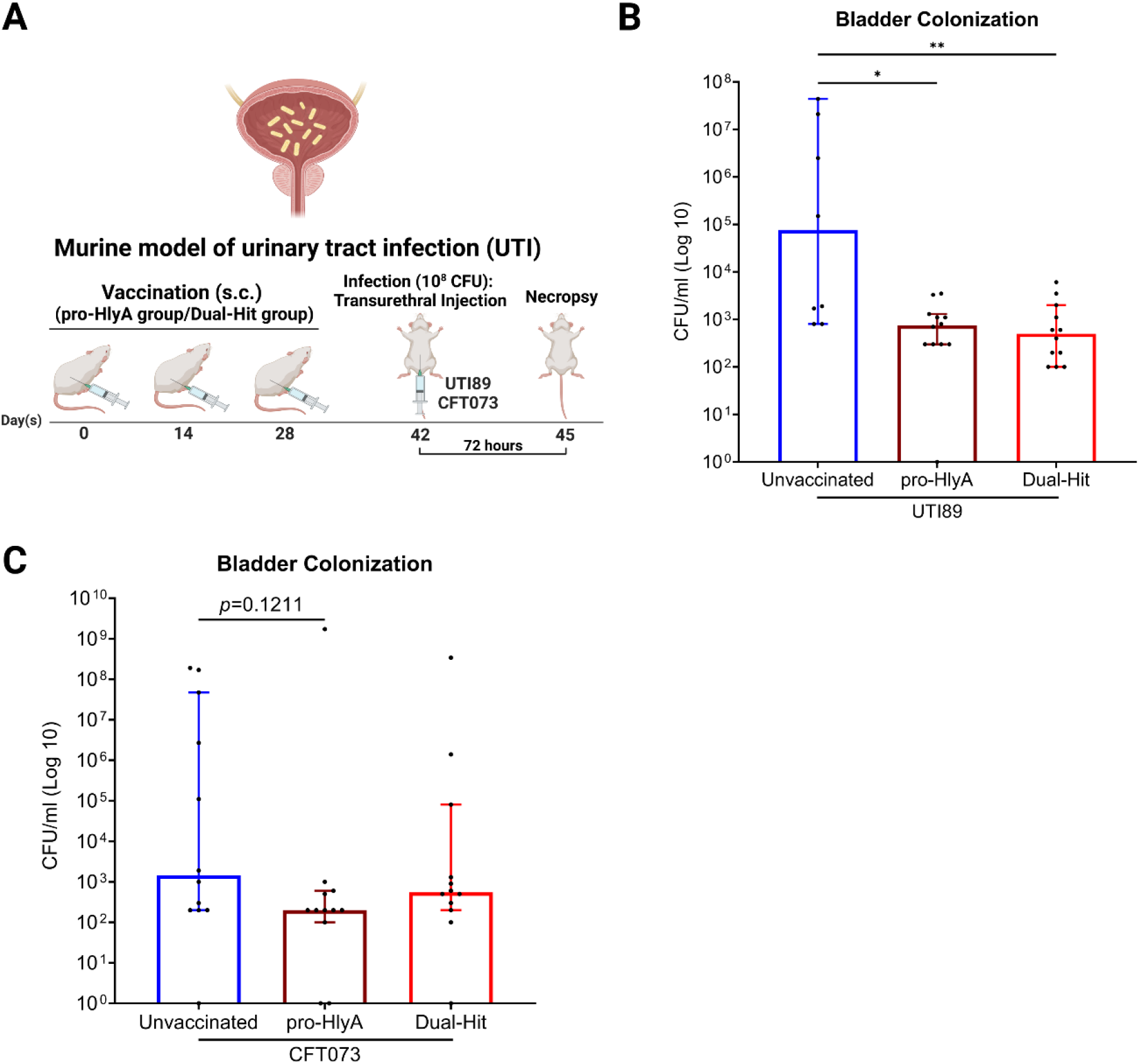
Evaluating the protective efficacy of pro-HlyA and Dual-Hit against UTI89 or CFT073 in th murine model of UTI. **(A)** The scheme of the murine UTI model using UTI89 or CFT073. Female BALB/cJ mice were subcutaneously immunized with either pro-HlyA (N=12/N=12), Dual-Hit (N=12/N=12) or remained unvaccinated (N=8/N=12), followed by a transurethral injection of 1 × 10^8^ CFU of UTI89 or CFT073. After 72 hours post-infection, bladders were harvested, homogenized, and plated to determine bacterial loads (CFU/ml). **(B)** Scatter plot with bar representing bladder UTI89 bacterial dissemination; **(C)** or scatter plot with bar representing bladder CFT073 bacterial dissemination.

For the experimental UPEC strain UTI89, our results demonstrated that both pro-HlyA and Dual Hit immunizations provided robust protection against UTI89 colonization in the bladder within the UTI model. In comparison to unvaccinated mice, the pro-HlyA vaccinated mice exhibited an approximate 2.01-log reduction in the median level of UTI89 in their bladder (Adjusted *P*-value, *P* = 0.0328). Similarly, the Dual-Hit vaccinated mice showed an approximate 2.18-log reduction in the median level of UTI89 (Adjusted *P*-value, *P* = 0.0094) (Fig. 7B). However, no significant differences were observed between the experimental groups and the control group for the experimental UPEC strain CFT073 (Fig. 7C).

### Immunization with Dual-Hit confers robust protection against a mixture of ten typical ExPEC strains in the murine model of mortality

In previous studies, we demonstrated the robust protective efficacy of Dual-Hit against UTI89 (ST95) and CFT073 (ST73) in murine models of bacteremia and mortality. Additionally, we found that immunization with pro-HlyA alone provided inadequate protection against the infections caused by ExPEC ST131 strains that lack the *hlyA* sequence in the murine mortality model. Consequently, we evaluated whether Dual-Hit could offer robust protective efficacy and significantly increase survival rates against a mixture of ten typical ExPEC strains (including ST95, ST73, and ST131) in the murine mortality model.

To assess the protective efficacy of the Dual-Hit against multiple sequence types of ExPEC strains in the murine mortality model, experiment group mice received three subcutaneous immunizations with Dual-Hit on days 0, 14, and 28, while a control group remained unvaccinated (Fig. 8A). On day 42, mice were intraperitoneally challenged with a mixture of ten typical ExPEC strains (1 × 10^8^ CFU/mouse in total), representing a range of common high virulent sequence types ExPEC strains (CFT073 (ST73), UTI89 (ST95), W0008 (ST127), JJ1886, JJ1901, JJ2050, JJ2528, JJ2547 (ST131), W0044 (ST405-like), and W0128 (ST648-like) in equal proportions). Over the next 10 days, mice were closely monitored for morbidity and mortality twice daily. Survival data were collected over time, and moribund or deceased mice were euthanized and necropsied to determine bacterial loads in kidneys, spleen, and liver. A small number of surviving vaccinated mice was randomly chosen and euthanized at 3 d.p.i, combined with the vaccinated mice that died within 1 d.p.i, forming the 3 d.p.i group, to assess changes in bacterial burden within infected organs over time (excluded from survival rate calculation). At 10 d.p.i all remaining vaccinated mice were euthanized, forming the 10 d.p.i group. Harvested organs were homogenized, and bacterial burdens within infected tissues were quantified by determining CFU.

**Fig 8.**
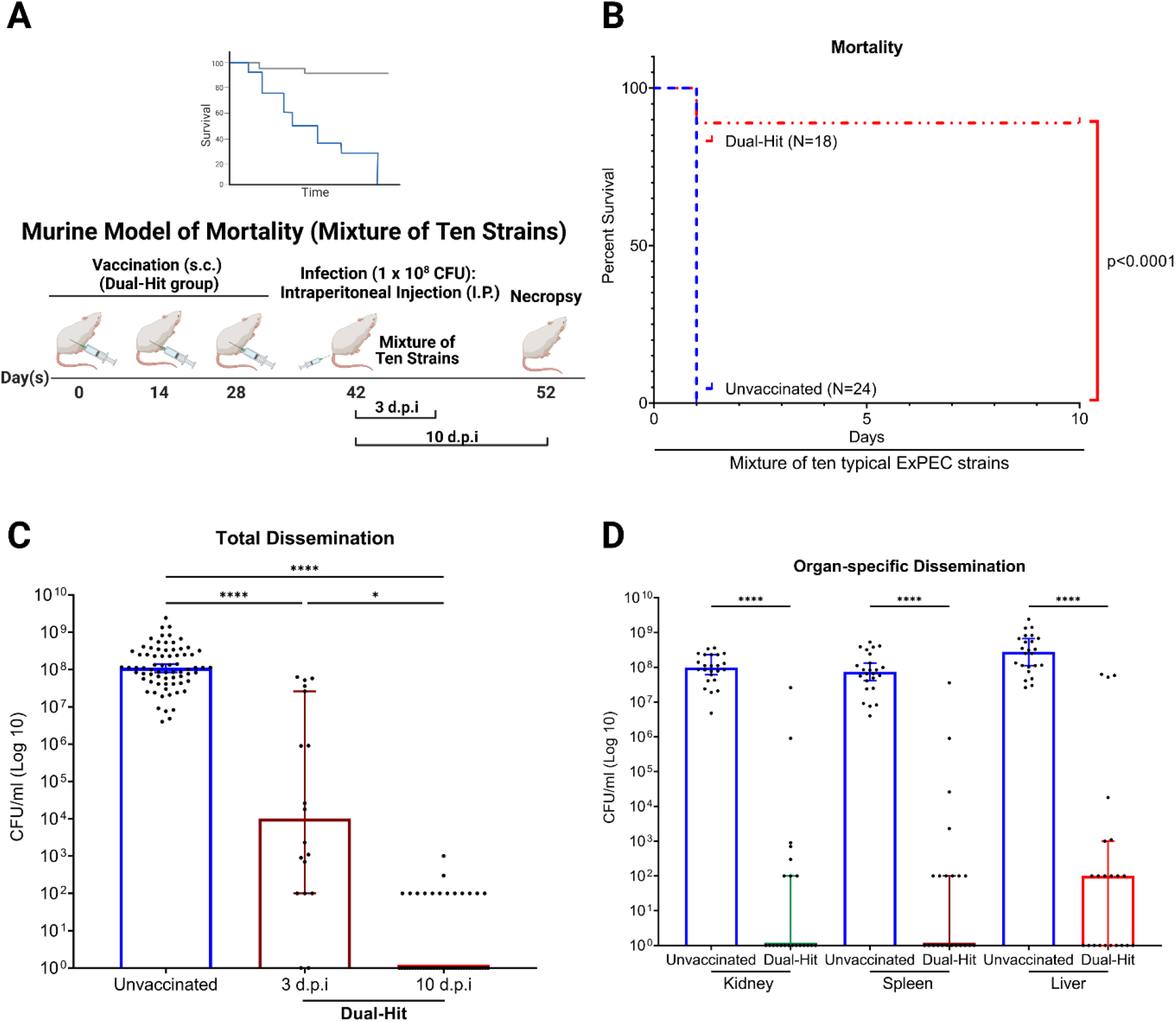
Evaluating the protective efficacy of Dual-Hit against a mixture of ten typical ExPEC strains infection in the murine model of mortality. **(A)** The scheme of the murine mortality model using a mixture of ten ExPEC strains. Female BALB/cJ mice were subcutaneously immunized with Dual-Hit (N=22), or left unvaccinated (N=24), followed by an intraperitoneal (I.P.) injection of 1 × 10^8^ CFU (in total) of a mixture of ten typical ExPEC strains. Mice were monitored twice daily for 10 days. The moribund or deceased mice were euthanized and necropsied to determine bacterial levels in organ (kidney, spleen, liver, CFU/ml). Vaccinated mice that died of infection within 1 d.p.i (N=2) and four randomly selected surviving vaccinated mice euthanized at 3 d.p.i (N=4) formed the 3 d.p.i group (N=6, excluded from survival rate calculation). All remaining surviving mice were euthanized at 10 d.p.i, forming the 10 d.p.i group (N=16). **(B)** Survival rates of Dual-Hit immunized mice following a mixture of ten typical ExPEC strains infection were analyzed using the Gehan-Breslow-Wilcoxon comparison. **(C)** Scatter plot with bar representing the total bacterial dissemination of mixture of ten typical ExPEC strains combining counts from all organs at 3 d.p.i and 10 d.p.i; **(D)** or the organ-specific bacterial dissemination of mixture of ten typical ExPEC strains in each organ type (combining counts from 3 d.p.i and 10 d.p.i) post-necropsy. Schemes were created in BioRender. Scatter plots with bars and Kaplan Meier survival curves were exported from Graphpad Prism 9 and annotated using BioRender.

Our findings revealed that unvaccinated mice died within 1 d.p.i, while 2 of the 18 Dual-Hit vaccinated mice died within 1 d.p.i during the 10-day observation period, resulting in a survival rate of 88.9% at 10 d.p.i (Adjusted *P*-value, *P* < 0.0001) (Fig. 8B). Bacterial burden measurements in Dual-Hit immunized mice corresponded with survival rate results. When combining counts from all organs, mice immunized with Dual-Hit demonstrated significantly reduced bacterial burdens at both 3 d.p.i (Adjusted *P*-value, *P* < 0.0001) and 10 d.p.i (Adjusted *P*-value, *P* < 0.0001) compared to unvaccinated mice. Compared to unvaccinated mice, the median bacterial burden in Dual-Hit vaccinated mice was approximately 4.04-log and 8.05-log lower at 3 d.p.i and 10 d.p.i, respectively (Fig. 8C). A 4.01-log reduction in the median level of ten ExPEC strains was also observed when comparing the bacterial levels in Dual-Hit vaccinated mice at 10 d.p.i to those at 3 d.p.i (Adjusted *P*-value, *P* = 0.0148), suggesting Dual-Hit immunization provided sustained and persistent protection against ExPEC infection over time (Fig. 8C). Furthermore, when combining the counts at both 3 d.p.i and 10 d.p.i and analyzing by each organ, Dual-Hit immunized mice demonstrated a significant reduction in bacterial loads across multiple organs compared to unvaccinated mice (Adjusted *P*-value, kidney, *P* < 0.0001; spleen, *P* < 0.0001; liver, *P* < 0.0001) (Fig. 8D). This reduction in bacterial burden was observed systemically across all collected organs, indicating a systemic reduction rather than localized effects conferred by Dual-Hit immunization.

## Discussion

ExPEC is the predominant cause of bacteremia and UTIs, persisting in both community environments and among hospitalized patients, leading to considerable hospitalization and mortality rates. The clinical management of ExPEC faces challenges, which are further exacerbated by the overprescription of antibiotics, the emergence of antibiotic-resistant ExPEC strains, and the global aging trend [21,92–93]. A vaccine targeting ExPEC represents a promising alternative strategy to address this issue, potentially mitigating the escalating global burden of antimicrobial resistance crisis and reducing hospitalization cost, thereby providing tremendous worldwide benefits.

In this study, we demonstrate that; (i) immunization with pro-HlyA or Dual-Hit elicited a rapid and robust protection against the highly virulent ExPEC strain UTI89 (ST95), reducing the bacterial burden of UTI89 in the murine bacteremia model; (ii) immunization with pro-HlyA or Dual-Hit increased survival rates following UTI89 infection, providing lasting and consistent protection in the murine model of mortality; (iii) immunization with pro-HlyA or Dual-Hit conferred partial protection against the highly virulent ExPEC strain CFT073 (ST73), decreasing the bacterial burden of CFT073 and increasing survival rates after CFT073 infection in the murine model of mortality; (iv) immunization with pro-HlyA or Dual-Hit reduced UTI89 induced cystitis in the murine model of UTI; (v) immunization with Dual-Hit significantly increased survival rates following infection by a mixture of ten typical ExPEC strains in the murine model of mortality, indicating the synergistic and broad-spectrum effects of the two antigens. Overall, our data indicate that both the inactive form of hemolysin, pro-HlyA, and Dual-Hit, a combination of the extracellular domains of the autotransporter SinH (SinH-3) and pro-HlyA, represent promising ExPEC vaccine candidates. We believe our findings offer an alternative research approach to the current ExPEC vaccine development efforts.

Hemolysin is a prevalent exotoxin produced by *E. coli* and significantly amplifies virulence in various clinical infections. Despite the relatively low abundance of hlyA in our phylogroup database as a whole, it is concentrated in highly virulent sequence types associated with ExPEC and UPEC infections, suggesting it plays a major role in these infections (Fig. 1A). Interestingly, the alignment and phylogenetic analysis of HlyA shows that the majority of instances of HlyA in what are generally considered intestine-associated phylogroups (A, B1, E) cluster together (Fig. 1B). This could suggest three different things: 1) the convergent evolution of a less-virulent (or more specialized) allele of hlyA, 2) a more promiscuous form of the pathogenicity island carrying hlyA, 3) increased horizontal transfer due to a higher likelihood of co-colonization. Given that the alleles found within non-B2 phylogroups are nested within the B2 phylogroup alleles, our results suggest that the B2 phylogroup acts as a reservoir for this virulence factor and that it occasionally spills over into other phylogroups, as we hypothesized previously [91].

Furthermore, while both pro-HlyA and Dual-Hit demonstrated high-efficiency protection against UTI89 in various murine models (bacteremia, mortality, and UTI), their protective efficacy against CFT073 in these models was not as robust as anticipated. The differential immunization protective efficacy against these two strains may stem from differences between UTI89 and CFT073. Although *E. coli* clones ST95 and ST73 frequently cause bloodstream infections and UTIs, a recent study revealed that UTI89 possesses a greater total number of genes that contribute to growth in urine and bladder colonization than CFT073. However, CFT073 appears to have more fitness factors than UTI89 [94,95]. Another study demonstrated that, while both UTI89 and CFT073 are clinical UPEC isolates that could cause infections for at least two weeks in similar proportions of mice, UTI89 infections could persist indefinitely, compared to the CFT073 infections began to clear two weeks after inoculation [96]. These findings suggest that

UTI89 might express more virulence factors on the bacterial surface compared to CFT073, leading to more persistent infections but also increasing detectability and bind-ability by vaccine specific antibodies against UTI89, leading to a higher protective efficacy of pro-HlyA and Dual Hit immunization.

Variations in immunization routes and adjuvant types significantly impact vaccine efficacy evaluation. Understanding these would enable optimization of the vaccine formulation and administration to maximize protective efficacy. For instance, intramuscular (I.M.) administration is the most used route for licensed vaccines and has been shown to elicit high immunogenicity in adult rabbits immunized with MecVax, producing antibodies against enterotoxigenic Escherichia coli (ETEC) H10407 and reducing intestinal colonization [99]. A clinical trial also demonstrated the safety and immunogenicity of the CS6-targeted candidate vaccine, CssBA, when administered intramuscularly [100]. Furthermore, both experimental and clinical evidence have indicated that mucosal immunization could efficiently induce local and distant systemic immune responses, as well as in the blood [97,98]. In previous studies, mice intranasally immunized with the iron receptor, FyuA, IutA, Hma, and IreA exhibited a robust and long-lived humoral immune response against UPEC challenge. Intranasal immunization with FyuA reduced UPEC strain 536 colonization following transurethral challenge, while IreA intranasally immunization significantly reduced CFT073 bacterial counts in the bladder [67,68]. Therefore, without compromising the robust systemic protection provided by pro-HlyA or Dual-Hit immunization, combining subcutaneous with either intramuscular or intranasal routes may present a more promising approach to enhance immune responses and improve protective efficacy of vaccinated mice against ExPEC infections in both blood (bacteremia) and mucosal (urinary tract).

The potential of a vaccine could also be significantly enhanced by formulating it with novel adjuvants, which effectively augment immune responses to the administered vaccine antigen [101]. In our study, we utilized aluminum salts (alum) as the adjuvant due to its proven safety. Alum is a clinically approved and widely used adjuvant in human vaccines, has been used for over 80 years in vaccine research and typically stimulates the Th2-type immune responses [102]. In a previous study, suitable adjuvants were screened for iron receptor-based immunization against UPEC infection, and they found that dmLT generated the most consistently robust antibody response in intranasally immunized mice, while Monophosphoryl-Lipid A (MPLA) and alum produced greater concentrations of antigen-specific IgG with intramuscular immunization [103]. This study suggests that dmLT, a mucosal adjuvant proven safe and potent through both preclinical and clinical studies, could be an alternative adjuvant in future research. Similarity, a recent study demonstrated that following bladder infection, highly T-helper type 2 (TH2)-skewed immune responses prioritized bladder epithelial repair after extensive exfoliation of epithelial cells, rather than bacterial clearance and even inhibition of Th1-mediated responses [104]. Therefore, MPLA is a potentially ideal adjuvant that could safely induce an appropriate level of Th1 response, enhancing Th1-mediated bacteria-clearing responses and increasing the ability to eliminate *E. coli* infection after vaccine immunization [105].

With the advancement of novel vaccine technology in recent years, we have also endeavored to develop an innovative mRNA vaccine targeting against ExPEC. Our mRNA vaccine encoded both *hlyA* and *sinH-3* sequences, incorporating an IL-2 signal peptide and a P2A self-cleavage sequence, built as the Dual-Hit mRNA construct, which is then encapsulated in cationic lipid nanoparticles. To evaluate the efficacy of the Dual-Hit mRNA vaccine against UTI89 in a murine mortality model, the experimental mice received three intramuscular (I.M.) immunizations with Dual-Hit mRNA vaccine (either low-dose or high-dose) on days 0, 14, and 28, while the control group mice were injected with an equal volume of Tris-buffer. On day 42, mice were intraperitoneally challenged with 1 × 10^8^ CFU/mouse UTI89 and monitored for morbidity and mortality and harvested organs as previously described (S2 Fig. A). Unfortunately, our results indicated that both control and low-dose group mice died within 1 d.p.i, while only 1 out of 7 high-dose vaccinated mice survived during the 10-day observation period. Furthermore, although one high-dose vaccinated mouse survived at 10 d.p.i, there is no statistical increase of both vaccinated groups in survival rate post-UTI89 infection (High-dose group, Adjusted *P* value, *P* = 0.1904), inferring the Dual-Hit mRNA vaccine only provided limited protection against UTI89 bacteriemia in murine mortality model (S2 Fig. B). Regarding bacterial burden measurements, combining counts from all organs, mice immunized with either low-dose (Adjusted *P*-value, *P* < 0.0001) or high-dose (Adjusted *P*-value, *P* = 0.0043) Dual-Hit mRNA vaccine had a statistically significant reduction in bacterial burdens compared to control mice. However, this level of protection was insufficient to increase the survival rate after UTI89 infection (S2 Fig. C). No discernable differences in organ-specific bacterial dissemination were observed among the three groups (S2 Fig. D). Although our Dual-Hit mRNA vaccine only provided limited protection against UTI89 bacteremia infection, the survival of 1 out of 7 high dose immunized mice at 10 d.p.i. highlights the potential of mRNA vaccines against bacterial infections. Insufficient immunization dosage may have contributed to these results, and future directions include increasing higher vaccination dosages and exploring higher protein expression efficacy via mRNA *in vivo*.

Antimicrobial resistance is a leading threat to global health currently. Developing the ExPEC vaccine presents a promising strategy to combat this growing global crisis and effectively reduce the incidence of antibiotic-resistant ExPEC infections to improve public health outcomes. In conclusion, our study demonstrates the promising protective efficacy of pro-HlyA and Dual-Hit immunizations against ExPEC infections in murine models. The observed reduction in bacterial burden and increased survival rates indicate the potential of these vaccine candidates for further development and evaluation. Furthermore, in this study, we bridged computational genomics and virulome vaccinology, attempted to block various steps of bacterial pathogenesis by synergizing multiple protein subunits associated with different virulence factors, and applied novel vaccine development technologies. We believe, our research contributes some innovative insights for future studies on vaccines against bacterial infections and offers a vaccine-development approach applicable to other severe MDR bacterial infections worldwide.

## Acknowledgments

We express our gratitude to James R. Johnson for granting us permission to utilize ExPEC strains JJ1886, JJ1901, JJ2050, JJ2528, and JJ2547 from his collection. The support for BCM Mass Spectrometry Proteomics Core is provided by the Dan L. Duncan Comprehensive Cancer Center NIH award and CPRIT Core Facility Award. Our appreciation goes to Professor Robert A. Britton, Professor Pedro A. Piedra, and Professor Pablo C. Okhuysen for their invaluable insights. Furthermore, we thank Ellen Vaughan, Carmen Gu Liu, Keiko Salazar, and Austen Terwilliger for engaging in discussions on specific virulence factors and Merry Wilks for her hard work on culture media supplying in this study.

## Notes

### Competing Interest Statement

The authors have declared no competing interest.

