## Supplementary material for "Progress toward the development of an effective vaccine for Extraintestinal pathogenic *E. coli* (ExPEC): The application of the multiple-protein subunits vaccine in different murine models": S1 Fig

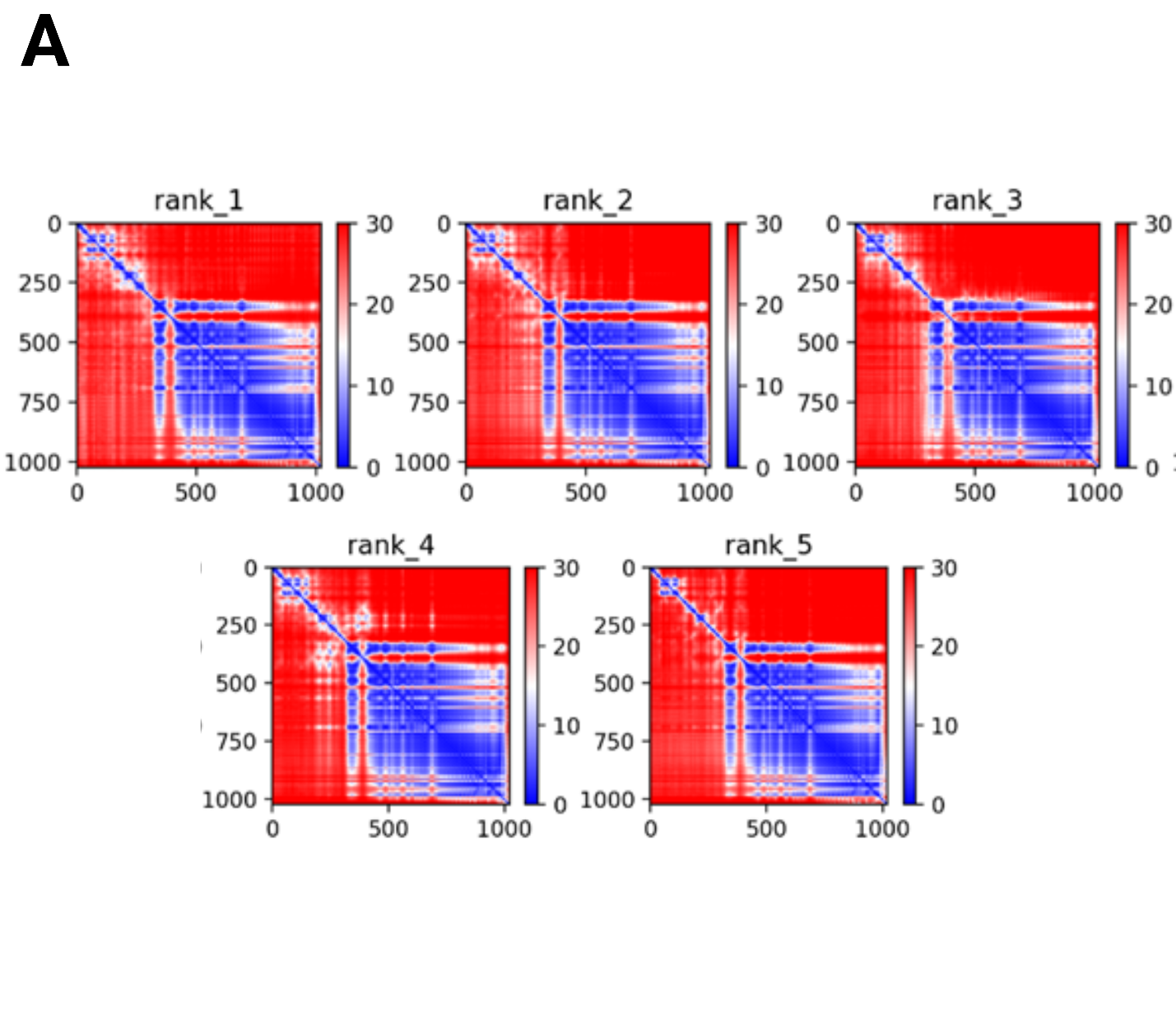

**S1 Fig.** **Predicted aligned error plots for five models used in predicting the structure of HlyA.** Each axis shows residue position, with position error between those residues depicted as heatmaps (Red: higher error, blue: lower error).
