## Supplementary material for "Progress toward the development of an effective vaccine for Extraintestinal pathogenic *E. coli* (ExPEC): The application of the multiple-protein subunits vaccine in different murine models": S2 Fig

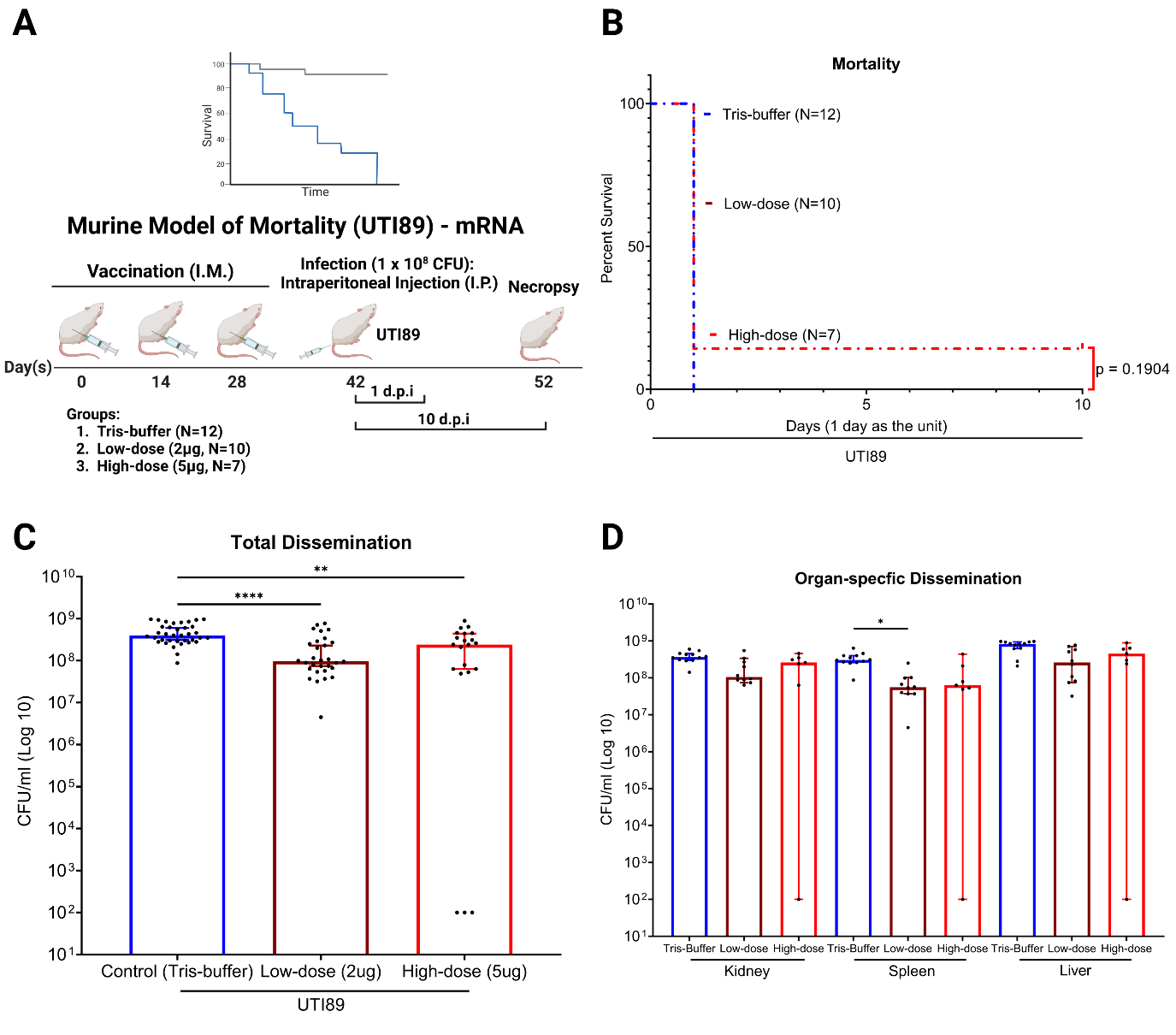

**S2 Fig. Evaluating the protective efficacy of Dual-Hit mRNA vaccine against UTI89 infection in the murine model of mortality. (A)** The scheme of the murine mortality model using UTI89. Male BALB/cJ mice were intramuscular (I.M.) immunized with Dual-Hit mRNA vaccine (low-dose, N=10; high-dose, N=7), or Tris-buffer (N=12), followed by an intraperitoneal (I.P.) injection of 1 × 10^8^ CFU (in total) of UTI89. Mice were monitored twice daily for 10 days. The moribund or deceased mice were euthanized and necropsied to determine bacterial levels in organs (kidney, spleen, liver, CFU/ml). **(B)** Survival rates of Dual-Hit mRNA vaccine immunized mice following UTI89 infection were analyzed using the Gehan-Breslow-Wilcoxon comparison. **(C)** Scatter plot with bar representing the total bacterial dissemination of UTI89 combining counts from all organs **(D)** or the organ-specific bacterial dissemination of UTI89 in each organ type post-necropsy. Schemes were created in BioRender. Scatter plots with bars and Kaplan Meier survival curves were exported from Graphpad Prism 9 and annotated using BioRender.
